# Optogenetic reduction of theta oscillations reveals that a single reliable time cell sequence is not required for working memory

**DOI:** 10.1101/2022.06.25.497592

**Authors:** Hyun Choong Yong, HaoRan Chang, Mark P. Brandon

## Abstract

In a delayed alternation spatial working memory task, hippocampal time cells fire during specific moments of the delay period to form a stable, sequential representation of the entire delay interval. The causal relationship between these sequences and working memory remains unclear. Similarly, hippocampal theta oscillations are thought to support working memory, primarily through the generation of time cell sequences. To causally examine these relationships, we optogenetically silenced the medial septal GABAergic theta-generating circuit during the delay portion of a delayed spatial alternation task. Without hippocampal theta oscillations, many time cells exhibited remapping, new time cells were recruited, and time cell information was increased; collectively resulting in a new time cell sequence during the delay period. Despite this remapping of time cells on random selection of theta-reduced trials, behavioral performance was unimpaired, demonstrating that working memory is not dependent on a single or unique time cell sequence during the delay period.

## INTRODUCTION

A neurobiological representation of *when* and *where* events occur is necessary for the expression of episodic memory (Tulving, 1983). Evidence from neurophysiological studies suggest that the spatial and temporal tuning of neurons in the hippocampus could serve as the basis for these aspects of cognition (Buzsáki & Moser, 2013; Eichenbaum, 2017). ‘Time cells’ exhibit temporally-tuned receptive fields that tile the delay period of a working memory task forming a ‘time cell sequence’, which might play a role in maintaining the contents of working memory during the delay (Kraus et al., 2013, 2015; MacDonald et al., 2011, 2013; MacDonald & Tonegawa, 2021; Pastalkova et al., 2008; Robinson et al., 2017; Sabariego et al., 2019; Salz et al., 2016; Shimbo et al., 2021; Wang et al., 2015). However, whether time cell sequences are necessary and/or sufficient to support working memory remains unclear. In prior work, time cells were observed while animals engaged in a memory task, and the fidelity of those time cell sequences correlated with memory performance (Gill et al., 2011; MacDonald et al., 2013; Pastalkova et al., 2008; Robinson et al., 2017; Wang et al., 2015). Importantly, time cell sequences also failed to emerge in a similar task that did not include a memory component (Pastalkova et al., 2008). Since these reports, however, time cell sequences have been observed in a ‘looping task’ with no memory demand (Mau et al., 2018), and do not emerge during the trace period of trace fear conditioning (Ahmed et al., 2020). Moreover, in one study, ablation of inputs from the medial entorhinal cortex spared hippocampal time cell sequences while memory performance was impaired (Sabariego et al., 2019). These reports thus suggest that time cell sequences could arise independently of working memory processes. Finally, studies that use direct and immediate optogenetic perturbations to the entorhinal-hippocampal circuit have reported concurrent disruptions of both time cell activity and memory (MacDonald & Tonegawa, 2021; Robinson et al., 2017). We suspect that these conflicting findings could be due to the methodologies used - lesion studies allow time for homeostatic regulation or other compensatory mechanisms (Maldonado et al., 2008; Otchy et al., 2015), while direct and immediate perturbation of hippocampal activity has the potential to influence a wide array of hippocampal and cortical processes, other than the activity of hippocampal time cell sequences (Li et al., 2019).

Beyond the search for the function of time cell sequences, computational models have generated clear hypotheses for how the hippocampus generates time cell sequences. Several popular models rely on hippocampal theta oscillations (6-10Hz), including models that use rhythmic persistent spiking and oscillatory interference (Hasselmo, 2008; Hasselmo & Stern, 2014), and/or theta-rhythmic inhibition to advance activity across the network (Haimerl et al., 2019; Wang et al., 2015). These models have received important experimental support from one study that used long-acting muscimol to silence the medial septum (MS), resulting in reduced theta power, disrupted time cell responses, and impaired memory performance (Wang et al., 2015). One important caveat to this study is that muscimol inhibits all medial septum cell types (GABAergic, cholinergic, and glutamatergic) which are hypothesized to support independent proposed mnemonic functions (Hasselmo, 1999; Roland et al., 2014). Further, these inactivations are maintained throughout an entire testing session - not specifically during the delay period. For these reasons, it has not been possible to attribute the perturbation in time cell sequences and the deficits in memory performance directly to the absence of theta oscillations and time cells during the delay portion of this task.

To untangle the complex relationship between theta oscillations, time cell sequences, and working memory, we used an optogenetic strategy to selectively silence the GABAergic population in the medial septum - a cell population necessary for the generation of theta oscillations in the hippocampus (Boyce et al., 2016). This approach allowed us to control the power of theta oscillations with similar efficiency to muscimol infusions selectively during the delay period in a delayed T-maze alternation task.

## RESULTS

### Theta oscillations are significantly reduced with temporal and spatial specificity

To investigate the contribution of theta oscillations to time cells and memory performance, we applied an optogenetic approach to specifically inhibit GABAergic neurons in the MS. First, the viral vector AAV2/DJ-Ef1α-Flex-ArchT-GFP was injected into the MS of VGAT::Cre mice to induce Cre-mediated expression of Archaerhodopsin (Han et al., 2011) in GABAergic neurons (n_ArchT_ = 14, n_Controls_ = 7). The expression was largely confined to the MS and the diagonal band of Broca (**Figures 1B and S1**). Next, microdrives with independently moveable tetrodes were implanted above and lowered into dorsal CA1 of the hippocampus (**Figures 1B and S1**). Mice were then trained in a delayed spatial alternation T-maze task to alternate between the left and the right arm and to run on a motorized treadmill (constant speed of 21.67cm/s) in the delay zone for 10 seconds between each alternation (**Figure 1A**). After passing a task performance criterion (80% correct for two consecutive days), the MS was inhibited on a random subset of 50% of trials during the delay period when mice ran on the treadmill. Single units and local field potential (LFP) recordings were simultaneously acquired from the pyramidal cell layer of dorsal CA1.

**Figure 1.**
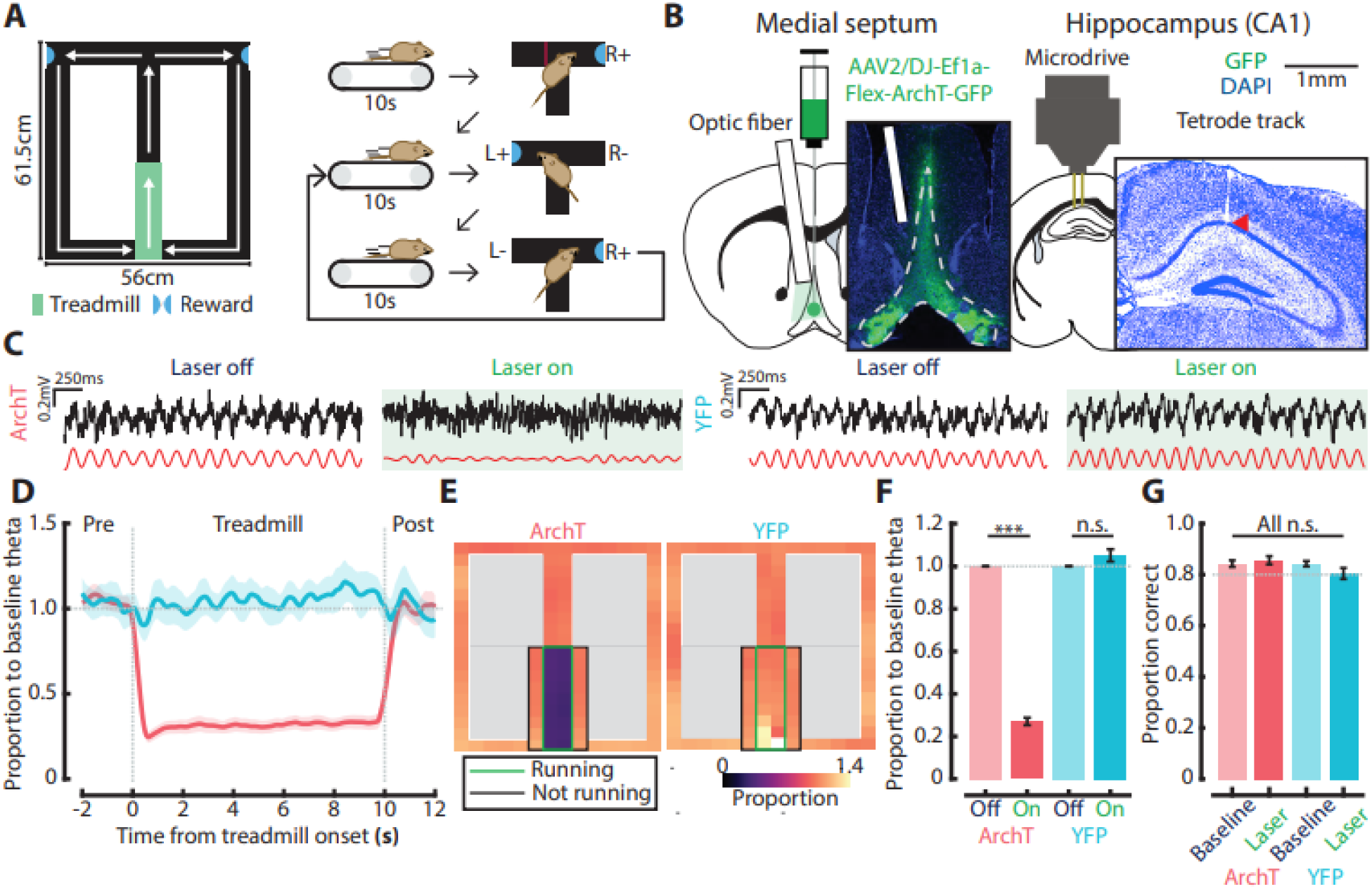
Theta oscillations (6-10Hz) are significantly reduced with temporal and spatial specificity during the delay period. (A) Schematic of the T-maze and overview of the delayed spatial alternation task. Mice were trained to alternate between the left and right arm of the maze while running on the treadmill (green) for 10 seconds between each alternation. Mice were rewarded with 0.02mL of water for a correct choice (+). (B) Histology of the MS showing virus expression (green), optic fiber location (white bar), and tetrode track tip in dorsal CA1 (red arrow). (C) Examples of raw (black) and filtered (red) LFP traces during laser-off and on trials in ArchT and YFP animals. (D) Proportion to baseline theta as a function of time. 2 seconds before and after the treadmill run are shown. Shaded area indicates the 95% confidence interval. Traces were smoothed over a 500ms window with a moving average. (E) Ratio between laser-on-theta and laser-off-theta as a function of location. Black square indicates the treadmill area while green square shows the proportion to baseline theta during the delay period. Left-going/returning, right-going/returning, and treadmill running periods were smoothed separately (S.D. = 2 bins) for visualization. (F) Proportion of laser-on-theta was significantly reduced compared to baseline theta in the ArchT group, but not in YFP controls. (Wilcoxon signed-rank test, ArchT: 0.269 (SEM: ± 0.017), p = 1.11 × 10^−9^ / YFP: 1.04 (SEM: ± 0.023), p = 0.16). (G) Behavioral performance between baseline and sessions with laser was not impaired in the ArchT group and YFP controls. (Wilcoxon signed-rank test, ArchT: baseline = 0.84 (SEM: ± 0.013), laser = 0.86 (SEM: ± 0.016), p = 0.46 / YFP: baseline = 0.84 (SEM: ± 0.011), laser = 0.80 (SEM: ± 0.022), p = 0.30). For all panels, n.s = non-significant, ***P < 0.001.

Hippocampal theta oscillations were substantially reduced when the laser was on in the ArchT group, but not in YFP controls (**Figure 1C**), consistent with a previous report during REM sleep (Boyce et al., 2016). Compared to laser-off trials, laser-on trials in the ArchT group had a 73±1.7% (SEM) reduction in theta power while YFP controls were unaffected (**Figure 1F**). To further investigate if the disruption was temporally and spatially specific to the delay period, the instantaneous theta power was obtained by continuous wavelet transform. To evaluate the temporal specificity of this effect, theta power was compared between laser-off and on trials before, during, and after the delay period (**Figure 1D**). Theta power in the ArchT group was reduced when the laser was turned on, and remained disrupted throughout the delay period before returning to baseline after the laser was turned off. On average, it took 202.26±5.81ms (SEM) to have theta power reduced to 50% of its baseline value. To interrogate if a similar level of specificity could be observed as a function of location, we obtained the occupancy-normalized theta power for individual bins, and compared between laser-off and on trials, separately for left-going/returning and right-going/returning trials (**Figure 1E**). Theta power was disrupted only when the ArchT group was running on the treadmill, but not when they were on the treadmill without running (i.e., waiting for the next trial to start or waiting for the door to open after the delay) or outside of the treadmill area. These results demonstrate that our approach allowed for strong control of the power of theta oscillations with high spatial and temporal precision while animals were performing the task.

### Intact behavioral performance despite partial remapping of time cells during optogenetic theta reduction

Having established that the reduction in theta oscillations was substantial and confined to the delay period, we investigated if animals’ memory performance was disrupted. Surprisingly, we did not observe any impairments in behavioral performance on the T-maze task (**Figure 1F**). This was the case even when the performance level was calculated separately for laser-off and on trials (**Figure S2B**). Moreover, behavioral performance was not correlated with the level of reduction in theta oscillations (**Figure S2A**).

We subsequently investigated whether temporal tuning of hippocampal CA1 neurons was affected by optogenetic manipulation. All analyses were carried out only using correct trials unless otherwise specified to ensure that provenance and destination are the same for each choice following the treadmill run. First, isolated single units were classified as a time cell when they passed the following criteria: (1) significant time information (bits/spike, (Skaggs et al., 1996)), (2) split-half reliability greater than the 95^th^ percentile of a shuffled distribution, (3) peak firing rate greater than 2Hz, and (4) overall mean firing rate less than 5Hz (to eliminate interneurons). The proportion of time cells identified within each laser condition was similar between the ArchT group and YFP controls (Pearson’s χ^2^ test of independence, ArchT off: 193 (38.8%), YFP off: 74 (31.4%), χ^2^ = 3.79, p = 0.10, ArchT on: 186 (37.4%), YFP on: 70 (29.7%), χ^2^ = 4.17, p = 0.082, corrected for multiple comparisons with Bonferroni).

Consistent with previous reports (Kraus et al., 2013, 2015; MacDonald et al., 2011, 2013; MacDonald & Tonegawa, 2021; Pastalkova et al., 2008; Robinson et al., 2017; Salz et al., 2016; Shimbo et al., 2021; Wang et al., 2015), time cells had firing fields at specific moments in time that collectively spanned the entire delay period (**Figures 2A-2B**). In response to optogenetic manipulation, partial remapping or ‘re-timing’ of time cells was consistently observed. The stability of the rate map between laser-off and on trials was significantly reduced in the ArchT group compared to YFP controls (**Figure 2C**). Unlike behavioral performance, the magnitude of time cell remapping was significantly correlated with the magnitude of theta reduction (**Figure S2C**). Furthermore, the time cell sequence for the ArchT group decorrelated faster over the course of the delay than YFP controls (**Figure 2D**). As another proxy measure of remapping, we compared the temporal peak firing location between laser-off and on trials. The conditional probability of the ArchT group had more dispersion from the diagonal compared to YFP controls, suggesting that time representations remapped between laser conditions (**Figure 2E**).

**Figure 2.**
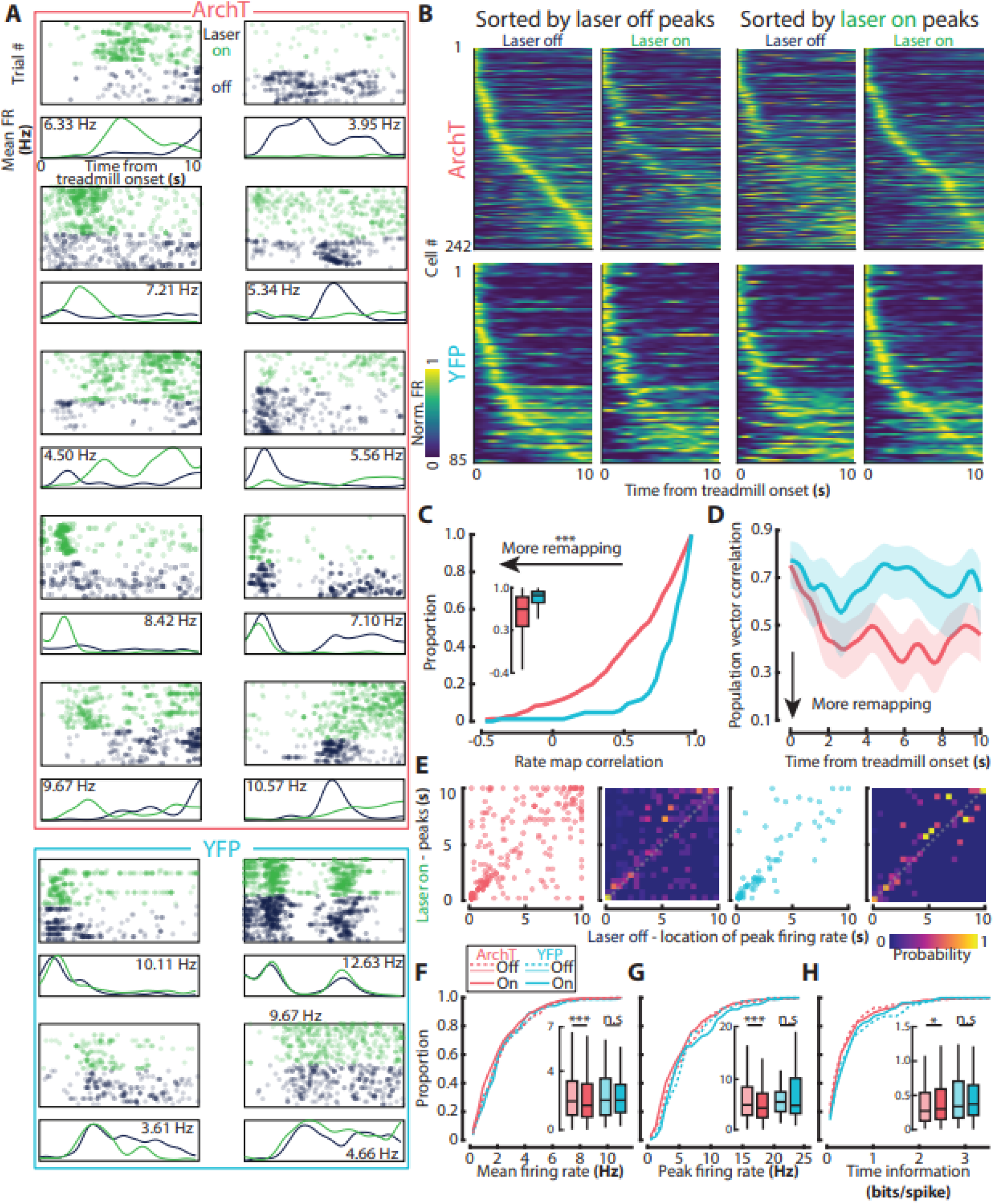
Time cells partially remap when theta oscillations are significantly reduced. (A) Individual time cell examples showing the effect of optogenetic inhibition of septal GABAergic neurons in ArchT (red box) and YFP (blue box) groups. Rasters for spike times, which are referenced to the beginning of each treadmill run, are shown with mean firing rate curves (laser-off: navy, laser-on: green). The firing rate shown in the box indicates the peak firing rate. (B) Sequence of time cells sorted by either their laser-off or on peaks. (C) Temporal rate map correlation coefficient for time cells between laser-off and laser-on conditions was reduced in the ArchT group (Wilcoxon rank-sum test, p = 4.36 × 10^−10^). (D) Population vector correlation coefficient of time cells (column-wise correlation in (B)) between laser-off and laser-on conditions over successive 200ms time bins (ArchT: red, YFP: blue). Two-sided confidence intervals were estimated using the procedure described in (Bonett & Wright, 2000). (E) Peak firing rate location during laser-on condition vs. peak firing rate location during laser-off condition (left) with conditional probability matrix (right). Peak firing rate location was more dispersed in the ArchT group compared to YFP controls, which has their peaks mostly concentrated along the diagonal in both conditions. (F) Mean firing rate of individual time cells between laser-off (dashed/shaded) and on (solid) conditions. In the ArchT group, there was a significant reduction in mean firing rate with the laser (Wilcoxon pairwise signed-rank test, p = 7.48 × 10^−7^), but not in YFP controls (p = 0.54). (G) Peak firing rate of individual time cells. Similar to mean firing rate, there was a significant reduction in the ArchT group with the laser (Wilcoxon pairwise signed-rank test, ArchT: p = 3.39 × 10^−4^, YFP: 0.33). (H) Time information of individual time cells. Significant increase in the ArchT group was observed (Wilcoxon pairwise signed-rank test, ArchT: p = 0.02, YFP: p = 0.88). Although there was a significant change in these time cell metrics, the overall shape of population distribution remained the same between laser-off and on conditions (two-sample Kolmogorov-Smirnov test, p > 0.05). For all panels, n.s = non-significant, *P < 0.05, ***P < 0.001.

Remapping of time cells was accompanied by significant changes in firing rates and time information when these properties were compared within cells between laser conditions (**Figures 2F-2H**). However, no differences were observed across laser conditions when the shape of population distribution was compared (CDF plots in **Figures 2F-2H**). These observations suggest that a new, partially remapped sequence emerges during laser-on trials. This indeed was the case when we sorted the sequence again by peak firing locations during laser-on trials (**Figure 2B**). Moreover, the proportion of time cells that were exclusively identified as a time cell in either laser-off or on trials was significantly higher in the ArchT group compared to YFP controls (Pearson’s χ^2^ test of independence, ArchT: 105/498 (21.1%), YFP: 26/236 (11.0%), χ^2^ = 11.07, p = 8.78 × 10^−4^). However, we did not observe a difference in the proportion of time cells in both laser-off and on trials (Pearson’s χ^2^ test of independence, ArchT: 137/498 (27.5%), YFP: 59/236(25.0%), χ^2^ = 0.52, p =0.47). Together, these results suggest that time cells remap when theta oscillations are disrupted, and a new sequence emerges to possibly support working memory.

### Temporal encoding is preserved by the emergence of a partially remapped sequence

To confirm that a partially remapped sequence emerges with reduced theta oscillations, we first investigated the split-half reliability between laser-off and on trials. This was done by randomly selecting a matched number of trials from laser-off and laser-on trials and calculating the Pearson correlation between resulting mean firing vectors. In line with our hypothesis, the split-half reliability between laser-off and laser-on trials in the ArchT group was significantly lower than other conditions (**Figure 3A**). Moreover, the split-half reliability between off-off and on-on trials did not significantly differ from each other, further suggesting that two representations reliably flip back and forth depending on the power of theta oscillations available for a given trial. Importantly, these changes were not observed in YFP controls (**Figure 3A**).

**Figure 3.**
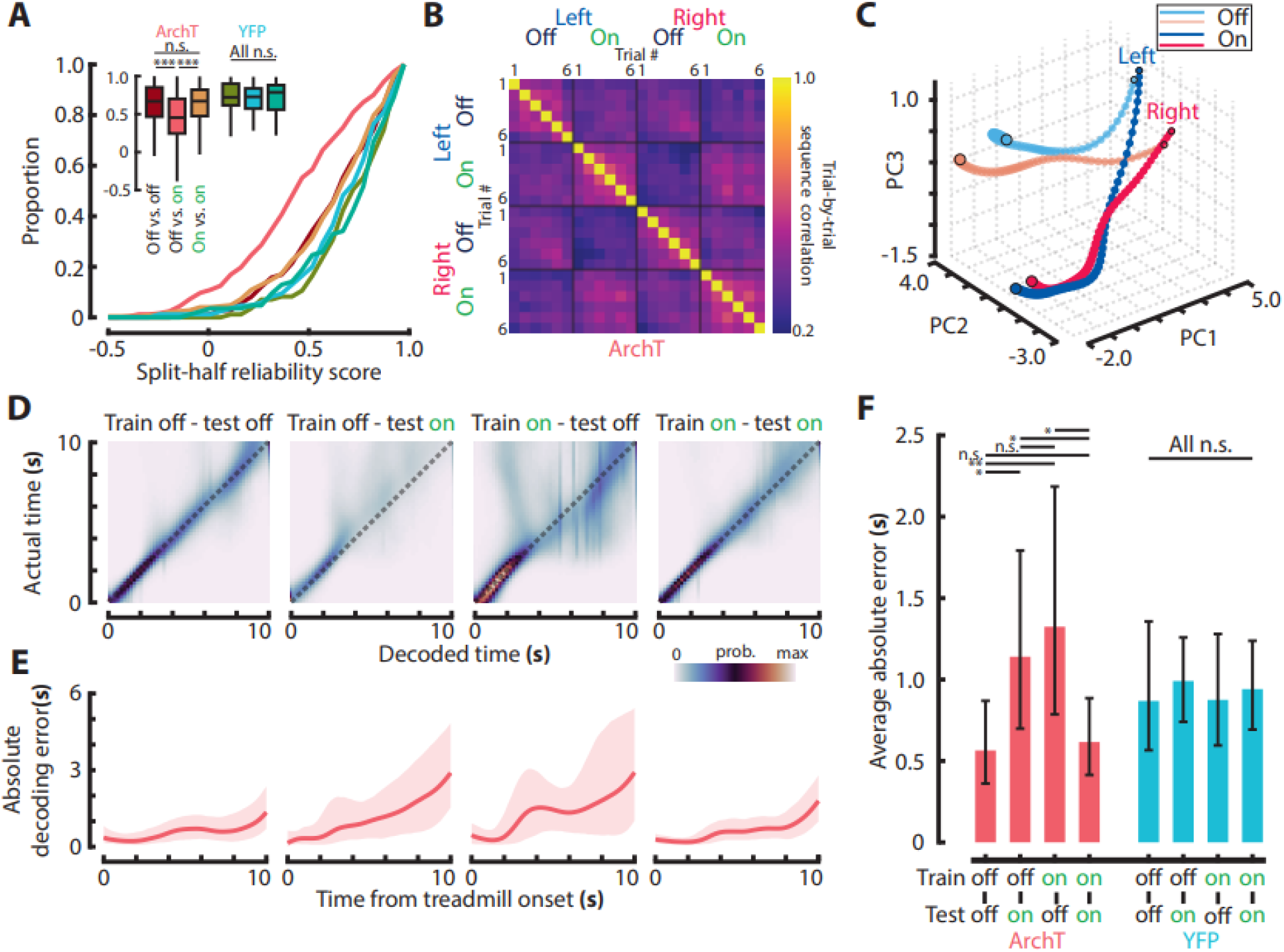
Temporal encoding is preserved during septal inactivation by the emergence of a partially distinct time code. (A) Split-half reliability for spike trains in the same condition (off-off, on-on), or across conditions (off-on). We randomly selected a matched number of trials from each condition and calculated the split-half reliability between resulting mean firing rate vectors. This was repeated 5000 times, and the mean of each null distribution was obtained. (Kruskal-Wallis test followed by post hoc tests, corrected for multiple comparisons with Bonferroni, ArchT: F(2, 723) = 55.11, p = 3.10 × 10^−12^, off-off vs. on-on: p = 0.99, off-off vs. off-on: p = 3.48 × 10^−10^, on-on vs. off-on: p = 4.39 × 10^−10^, YFP: F(2, 252) = 1.38, p = 0.50). Reduced split-half reliability in off-on condition suggests the emergence of remapped time code when theta oscillations are significantly reduced. (B) Trial-by-trial sequence similarity correlation matrix. Each pixel represents a correlation of sequences between a pair of trials. Note the checkerboard pattern, indicating remapping between laser-off and on conditions. Only those time cells with at least 6 trials in all four conditions were included in this and all subsequent analyses in this figure (ArchT = 178/242, YFP = 69/85). (C) 3D projection of neural trajectories in all four conditions (red = left, blue = right, graded color = laser-off, solid color = laser-on). The dot size indicates the time from the treadmill onset with larger ones indicating later in time. Black circles mark the beginning and end of the treadmill run. Axes are three first principal components with arbitrary units. This was plotted for visualization purposes only. (D) Confusion matrices of maximum likelihood decoding of time (see methods). (E) Absolute decoding error at each time bin across training conditions. Shaded area is 95% confidence interval. (F) Average absolute decoding error across training conditions in ArchT (red) and YFP (blue). Mean and 95% confidence interval for each training-testing condition are shown. Multiple paired bootstrap sample tests for equal mean absolute error were used to test the significance across different training conditions within ArchT and YFP groups. P-values were Bonferroni-corrected for multiple comparisons. Refer to Table S2 for p-values. For all panels, n.s = non-significant, *P < 0.05, **P < 0.01, ***P < 0.001.

Next, we visualized if a sequence in each trial was reliable within the same laser condition, but different between laser conditions. When visualizing the trial-by-trial sequence for the ArchT group, we observed a checkered pattern, suggestive of a reliable sequence representation within the same laser condition and decorrelated sequence representations between laser conditions (**Figures 3B-S3B**). This pattern was observed regardless of the minimum number of trials needed to be included in the analysis (**Figure S3C**). Indeed, Principal Component Analysis (PCA) shows that neural trajectories take markedly different paths depending on the laser condition (**Figure 3C**), but this effect was not observed in YFP controls (**Figure S3**).

To further confirm that a partially remapped sequence emerges between laser conditions, we utilized a maximum likelihood estimator to decode time of treadmill delay based on the population activity. In the ArchT group, the absolute decoding errors were comparable between laser-off and on conditions with a mean of ∼0.5s (where the theoretical chance level would be 3.33s) (**Figures 3D-F**). However, when the estimator, trained on the neural activities during laser-off trials, was used to decode laser-on trials (or *vice versa*), the absolute error increased substantially. In contrast, the levels of decoding error were maintained across training-testing conditions for YFP controls (**Figures 3F-S4A**). These results were consistent regardless of the number of cells and the minimum number of trials that were included in the training of the decoder (**Figures S4B-C**). These outcomes further outline the view that an altered, albeit similarly accurate, time sequence emerges with the reduction of theta oscillations.

### Trajectory-dependent time cells were resilient to optogenetic theta reduction

Given that behavioral performance was not disrupted following significant theta reduction and partial remapping between laser conditions, we hypothesized that trajectory-dependent time cells, signaling the past or future trajectories (Frank et al., 2000; Kinsky et al., 2020; Pastalkova et al., 2008; Wood et al., 2000) of animals, would remain stable despite this perturbation. We reasoned that this subset of time cells may be sufficient to guide behavior when other time cells remap during reduced theta oscillations. Time cells were identified to be significantly modulated by the trajectory when the difference in mean firing rate between left-going and right-going trials was greater than the 95th percentile of a shuffled distribution (Kinsky et al., 2020). This resulted in 46 and 12 trajectory-dependent neurons in the ArchT group and YFP controls, respectively. The proportion of trajectory-dependent neurons did not differ between the ArchT group (19%) and YFP controls (14%) (Pearson’s χ^2^ test of independence, χ^2^ = 1.03, p = 0.31) (**Figure 4A**).

**Figure 4.**
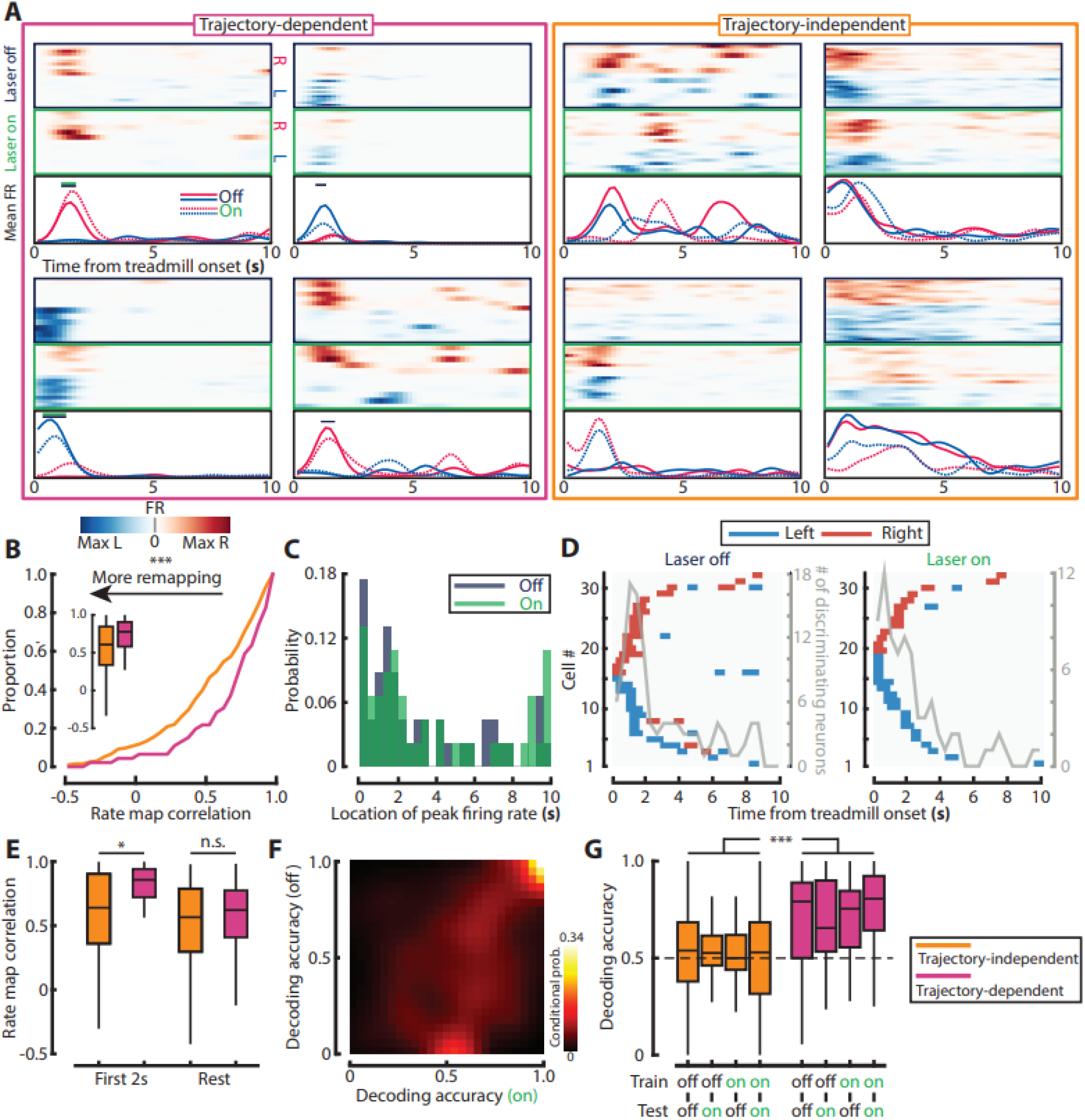
Trajectory-dependent time cells maintain more stable representations than trajectory-independent time cells. (A) Examples of trajectory-dependent (pink box) and trajectory-independent cells (orange box). Trial-by-trial firing rate is shown as a function of time, sorted by their choice following a treadmill run (left: blue, right: red) and the laser condition (off: blue box, on: green box). Mean firing rate is shown at the bottom for each condition (off: solid, on: dashed, same color scheme as described above). Solid horizontal line on top of the mean firing rate curve indicates significantly discriminating fields (off: blue, on: green) (B) Temporal rate map correlation coefficient, comparing trajectory-dependent and trajectory-independent cells. (Wilcoxon rank-sum test, ArchT: p = 0.006, YFP: p = 0.79). (C) Location of peak firing rate during laser-off (dark blue) and on (green) trials for trajectory-dependent cells. (D) Location of significantly discriminating fields for each trajectory-dependent cells during laser-off and on. Color scheme is the same as individual examples. The y-axis on the right indicates the number of trajectory-dependent cells that have a significantly discriminating field for a given time period. Note that location of peak firing rate and significantly discriminating fields are largely within the first 2 seconds of the delay period. (E) Temporal rate map correlation, comparing trajectory-dependent (pink) and trajectory-independent cells (orange) with their peak firing rate within the first 2 seconds or after. (Wilcoxon rank-sum test, first 2 seconds: ArchT: p = 0.015, YFP: p = 0.43, after 2 seconds: ArchT: p = 0.52, YFP: p = 0.97). (F) Conditional probability matrix showing the accuracy of a decoder built to predict the animals’ subsequent choice (left or right) based on firing patterns during the delay period. Note the diagonal in the upper right part of the matrix, indicating the consistent accuracy across laser-off and on conditions. (G) Decoding accuracy of choice (left or right) for trajectory-dependent and trajectory-independent cells (Robust 2-way mixed bootstrapped ANOVA with 20% trimmed means, number of bootstrap: 5000, only main effect of cell identity (trajectory-dependent vs. trajectory-independent) was significant (p < 1.00 × 10^−4^). Refer to Table S3 for ANOVA table, and Figure S5/ Table S3 for data from YFP controls. For all panels, n.s = non-significant, *P < 0.05, ***P < 0.001.

Trajectory-dependent time cells showed greater rate map stability compared to trajectory-independent time cells (**Figure 4B)**. Similar to a previous report (Pastalkova et al., 2008), we observed that trajectory-dependent time cells were highly concentrated towards the beginning of the treadmill run (**Figures 4C-D**). Indeed, according to our neural trajectory visualization and time decoding analysis (**Figure 3C-E**), decoding error at the beginning was lower compared to later portions of the delay. Thus, we separated time cells by their peak firing location during laser-off trials and compared the rate map correlation between trajectory-dependent time cells and trajectory-independent time cells. This revealed that trajectory-dependent time cells were more stable than trajectory-independent time cells during the first 2 seconds on the treadmill, while the stability of trajectory-dependent and trajectory-independent cell groups did not differ in the remainder of the treadmill delay period (**Figure 4E**).

To further corroborate these findings, we designed a decoder to predict the animals’ subsequent choice based on spiking activities during the delay period (see Methods). The decoder was first trained to discriminate between left-going and right-going trials during laser-off and on trials separately, and prediction accuracy was obtained by leave-one-out cross-validation. Under both conditions, cells that expressed high decoding accuracy during laser-off trials maintained their accuracy during laser-on trials (**Figure 4F**). To examine if the decoding accuracy was stable across laser conditions, we trained the decoder on laser-off trials and tested the accuracy for decoding laser-on trials, and vice versa. Across train-test conditions, trajectory-dependent time cells had higher decoding accuracy compared to trajectory-independent time cells, while the same level of accuracy was maintained across train-test conditions (**Figure 4G**). These results suggest that a small population of cells that are more relevant to behavioral performance maintain stable time representation despite strong theta reduction, which could be sufficient to support working memory.

## Discussion

To disentangle the relationship between theta oscillations, time cell sequences, and working memory, we optogenetically suppressed GABAergic medial septal neurons exclusively during the delay period of a delayed alternation memory task. Surprisingly, we found that strong suppression of theta oscillations did not impair working memory despite leading to remapping and the emergence of a distinct time cell sequence during the delay. Our results suggest that, in the absence of theta oscillations, working memory could be supported by intact trajectory-dependent time cells at the beginning of the delay period that signal behaviorally-relevant past and future trajectories. Taken together, these results suggest that theta oscillations are not required during the delay period of a delayed spatial alternation task. While theta oscillations may play a key role in maintaining stable trial-by-trial time cell sequence, a single, task-persistent time cell sequence is not required for working memory.

Our results also speak to the putative mechanisms that support time cell sequences. We demonstrate that the presence of theta oscillations allows for a consistent readout, across trials, of a single sequence. Without theta oscillations, a new sequence emerges, particularly evident after ∼2 seconds on the treadmill. The stability of time cells at the beginning of the treadmill may be driven by immediate prior experience and sensory inputs, while expression of the remainder of the sequence - when sensory information is no longer informative - is dependent on an internally driven theta-dependent process, similar to mechanisms proposed in time cell models (Haimerl et al., 2019; Hasselmo & Stern, 2014; Wang et al., 2015). Without this theta-dependent mechanism, retrieval of the original sequence fails and is replaced by a new sequence.

We speculate that this new sequence could emerge due to a shift in temporal information arriving to CA1 from CA3 and the medial entorhinal cortex (MEC). It was previously reported that both CA3 and the MEC contain time cells (Kraus et al., 2015; Salz et al., 2016). Inhibition of MEC input to the hippocampus has been shown to induce remapping of time cells (Robinson et al., 2017), and time cell sequences have been observed to remain intact following extensive lesions to the MEC (Sabariego et al., 2019). These studies strongly suggest that CA3 provides CA1 with sufficient temporal information to express a time cell sequence. An important parallel can be drawn with works from the spatial coding literature: neither lesions to CA3 or MEC prevent spatial coding in CA1 (Brun et al., 2002, 2008), demonstrating that either input is sufficient to drive spatial coding in this subregion. In septal inactivation studies, which disrupt the tuning of MEC grid cells (Brandon et al., 2011; Koenig et al., 2011), spatial coding in both CA1 and CA3 remains intact (Brandon et al., 2014; Koenig et al., 2011; Wang et al., 2015). Given that grid cells can function as time cells in the delayed T-maze task, it is likely that these cells are similarly disrupted in the temporal, or distance traveled, domain (Kraus et al., 2015) during optogenetic silencing of septal GABAergic populations. Thus, the remapping reported here may result from a disruption in the integration of parallel streams of temporal information arriving from the MEC and CA3, whereby only CA3 temporal inputs remain intact and can drive a new sequence in CA1 without the presence of theta oscillations. Indeed, CA3 is a strong candidate for retrieval of temporal sequences in the absence of changing sensory information due to its recurrent, autoassociative connectivity (Marr, 1971; McNaughton & Morris, 1987). Future experiments should confirm if MS inactivation disrupts time cells in MEC.

Our results are the first to demonstrate that switching between partially remapped time cell sequences does not impact working memory, which is a markedly different result from prior reports that used other circuit optogenetic manipulations to degrade time information and memory. In previous work, transient inactivation of the MEC disrupted CA1 time cell sequences substantially, and induced memory impairments (Robinson et al., 2017). In this case, time cell information, time cell sequences and behavioral performance were disrupted in both laser-on and laser-off trials, suggesting that direct manipulations of the MEC had a relatively long impact on entorhinal-hippocampal physiology that persisted across trials. Moreover, transient CA2 inactivation did not appear to recruit a new time cell sequence in prior work, and in contrast to that reported here, time cells exhibited increased firing rates and decreased time information (MacDonald & Tonegawa, 2021). Similarly, full medial septum inactivation with muscimol led to both decreased time field information and memory (Wang et al., 2015). It is noteworthy to add that these behavioral impairments may have also been the consequence of non-selective inhibition of all septal neurons, as recent studies highlight the importance of non-GABAergic septal neurons for working memory performance. Stimulation of cholinergic neurons during the delay (Y. Zhang et al., 2021), and inhibition or random rhythmic perturbation (Etter et al., 2021; Gemzik et al., 2021) of all septal cell types during the delay period led to behavioral impairments. In our study, despite switching to a partially remapped time cell sequence, selective inactivation of septal GABAergic neurons led to decreased firing rates and *increased* time information. In contrast to other reports, the maintenance of high temporal information during septal GABAergic inactivation may help to explain why working memory remained intact.

While the baseline and theta-reduced time cell sequences were statistically different, these sequences were not orthogonal. Trajectory-dependent time cells in the first 2 seconds remained stable (**Figure 4E**), and the population correlation following the initial 2 seconds was maintained around 0.4 (**Figure 2D**). Thus, at any given moment in time, the neural trajectory at the population level could be similar enough for downstream readers to generate correct tuning responses in the delayed spatial alternation task.

An important alternative perspective, given that partially remapped time cell sequences were present both during baseline and theta-reduced trials with high temporal information, is that animals were capable of using both sequences to perform the task. Similar findings have been reported on the time scale of days and weeks, whereby behavioral performance is maintained despite substantial representational drift (Driscoll et al., 2017; Levy et al., 2021; Ziv et al., 2013). In the current study, representational content was immediately altered optogenetically, yet the hippocampal representation during the delay was sufficient to support behavioral performance. This view suggests that downstream readers important for behavioral decisions can rapidly switch to interpret distinct temporal sequences that subserve the same behavioral outcome.

## Materials and Methods

### Subjects

VGAT-ires-Cre male mice (the Jackson Laboratory, stock #01692) were housed individually on a 12-h light/dark cycle. Only male mice were used in this experiment, as female VGAT-ires-Cre mice were considerably smaller (**Table S4**) and have increased difficulty to support the weight of electrophysiology preamplifier, tether, and optic patch cord. All experiments were carried out during the light cycle, and all experiment procedures were approved by McGill University and Douglas Hospital Research Centre Animal Use and Care Committee (protocol #2015-7725) and performed following Canadian Institutes of Health Research Guidelines. The previous work has demonstrated the selectivity of this transgenic mouse line for GABAergic neurons in the medial septum (Boyce et al., 2016).

### Surgeries

Prior to surgery, mice were initially anesthetized with isoflurane (5% at 1.5% oxygen) and injected subcutaneously with carprofen (0.01mg/g) and 1mL of sterile saline. Mice were maintained at 0.5%-2.5% isoflurane with 1.5% oxygen throughout the surgery. Eye lubricant and a heating pad were used to keep eyes hydrated and maintain the body temperature, respectively. All surgeries were carried out on a stereotaxic frame (David Kopf Instruments, Inc) and prior to surgeries, all surgical tools were sterilized with a glass bead sterilizer. Two surgeries per mouse (viral injection followed two to four weeks later by a combined optic fiber/microdrive implantation) were needed to prepare them for recording.

Male mice at 8-12 weeks of age were injected with 600nL of AAV2/DJ-Ef1α-Flex-ArchT-GFP (1.1×10^13^) or AAV2/DJ-Ef1α-Flex-GFP (8.2×10^12^), obtained from the Neurophotonics Centre at the University of Laval. Mice received a single injection at AP: +0.82 / ML: 0.00 / DV: -4.80 with a glass pipette connected to Nanoject II injector (Drummond Scientific, Inc) at a flow rate of 23nL/s, or at AP: +0.82 / ML: -0.5 / DV: -4.8 (angled at 4.5° towards the midline) with a 28G cannula connected to a pump (Harvard Apparatus). Glass pipette or cannula was retracted after waiting for 10 minutes following the injection.

Two to four weeks following the injection surgery, an optic fiber (CF230, Thorlabs; implant site: AP: +0.82 / ML: -0.5/ DV: -3.64 at 4.5°) and a microdrive (Versa drive, Axona Ltd; implant site: AP: -1.8 / ML: +1.6) were implanted above the medial septum and dorsal CA1, respectively. Prior to implant surgery, tetrodes were gold-plated using the NanoZ (Neuralynx) to lower the impedance below 250kΩ at 1004Hz. Two stainless anchor screws (B000FN0J58, Antrin Online) were placed in front of the inferior cerebral vein, and the optic fiber was secured to the skull with dental cement (Patterson Dental, Inc). After dental cement was dry, two more anchor screws were placed on the contralateral hemisphere. A third craniotomy was made on top of the cerebellum to place a ground screw. After putting the microdrive and the ground screw in place, both holes were initially secured with a silicone adhesive (Kwik-Sil, World Precision Instruments) and then covered with dental cement. Once the cement was dry, tetrodes were lowered 250um below the surface of the brain.

Mice were maintained on the heating pad and monitored until they fully recovered after each surgery. For the first seven days after each surgery, mice were given a soft diet along with carprofen gel (MediGel CPF, ∼5 mg kg−1 each day) as an analgesia and a diet boost (DietGel Boost).

### Behavioral training on the delayed alternation with treadmill T-maze task

At least seven days following the implant surgery, mice were placed on water restriction and their weight was maintained above the 85% of their *ad libitum* weight throughout the experiment. On the first day, mice were habituated to the T-maze (MazeEngineers, Inc) by allowing them to freely explore the T-maze for 30 minutes. Water droplets were placed throughout the maze except for the treadmill area to encourage explorative behavior. Mice were deemed to have passed the habituation when they voluntarily explored the entire maze within 30 minutes. Following the habituation, mice were habituated to run on the treadmill. In each treadmill habituation session, the speed of the treadmill was held constant, but the duration of treadmill run was increased from 5s to 15s in 5s increment after every 5 trials (total of 15 trials). The speed was gradually increased from 8.33cm/s to 21.67cm/s with an increment of 3.33cm/s between sessions (5 sessions in total). Following the treadmill training, mice were trained to drink from reward ports which were located at the end of each choice arm. Reward ports were loaded with 0.02mL water and mice were given two minutes to drink from the port. After two minutes, mice were manually transferred to the opposite reward port and given another two minutes to drink. Mice were deemed to have familiarized themselves with the reward ports if they drank 8 out of 10 times and two skipped trials were not from the same side.

Once mice passed all pre-trainings, they were trained to perform a delayed spatial alternation task with a delay of 10s between each alternation. During this 10s delay, mice ran on the motorized treadmill at 21.67cm/s for the entire delay period. Each session started with a 10s of treadmill run and then a forced trial (forced to go to the left or right with the door to the opposite arm closed). The forced trial was pseudo-randomly selected. After retrieving the water reward of 0.02mL from the reward port, mice had to come back to the treadmill to initiate the second trial. From the second trial, mice were able to choose between the left and right arm, however, in order to receive the water reward, they had to choose the arm opposite to the arm they had previously visited. If an incorrect choice was made, mice were placed in a correction trial with the same 10s run on the motorized treadmill until they made a correct choice. Mice were considered to have learned the task if they had 80% correct with more than 30 trials (excluding the forced trial) in 30 minutes for two consecutive sessions.

### Optogenetic inhibition of medial septal GABAergic neurons

Once mice reached the criteria, a green laser (520nm wavelength, maximum power output: 60mW, Doric Lenses) was pseudorandomly used to inhibit GABAergic neurons in the MS. In a closed-loop system, the light was turned on in 50% of trials, and triggered on and off by the treadmill start and end TTL, respectively. The estimated light power at the tip of the optic fiber was never allowed to exceed 20mW.

### In vivo electrophysiological recordings

Both spikes and local field potential recordings were sampled and digitized at 32kHz using a Digital Lynx recording system (Neuralynx, Inc). Signals were amplified and band-pass filtered between 0.6kHz and 6KHz. Threshold was adjusted prior to each recording, ranging from (30 µV to 65 µV). Sleep recordings were used to guide tetrodes to the CA1 region of the hippocampus while mice were resting in their home cage. Both Neuralynx and Open Ephys systems were used for this purpose.

### Analysis of theta power

The power spectrum for local field potentials was obtained using multitaper method included in the Chronux toolbox (*mtspectrumc* with NW = 3 and K = 5; (Mitra & Bokil, 2007). Theta power was calculated by taking the area within 1Hz of the maximum power in the theta range (6-10Hz). Baseline theta power was obtained by taking the average of theta power during laser-off trials, and mean reduction in theta power for each session was obtained by dividing each laser-on trial with baseline theta power and averaging across laser-on trials. For each channel, the theta-delta ratio was obtained by taking the ratio between mean theta power (6-10Hz) and mean delta power (2-4Hz) during laser-off segments. A channel with the highest theta-delta ratio was used in the LFP analysis.

Instantaneous theta power was estimated with the MATLAB built-in function *cwt* using the Morlet wavelet. This was used to calculate the reduction in theta power as a function of time and location. For time, mean reduction in theta power before, during, and after the treadmill run was compared as described above. For location, T-maze was linearized, and x-y coordinates, which are sampled at 30Hz, were linearly interpolated to match the LFP sampling rate (500Hz). After binning theta power, mean reduction in theta power was calculated in the same way as described above. Time points falling into the cone of influence were excluded from these analyses.

### Identification of time cells

Spike times on the treadmill were first referenced to the beginning of each treadmill run. Spike times were binned into 200ms bins, and convolved with a Gaussian kernel (S.D. = 400ms). The smoothed spike trains were used throughout the analysis. Time information *I* was derived for individual neurons as previously described (Skaggs et al., 1996):

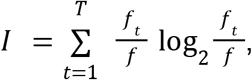

To assess the stability of the representations of neurons across trials, a reliability score was obtained by constructing the mean firing rate vectors out of even and odd trials, and by computing the Pearson correlation coefficient between the two vectors. Neurons need to express time information and split-half reliability higher than the 95^th^ percentile of their shuffled distribution. To ensure that cells are active on the treadmill, peak firing rate and mean firing rate had to be greater than 2Hz and 0.5Hz, respectively. Any cells with overall mean firing rate of equal to or greater than 5Hz were excluded from the analysis. Cells had to pass these criteria in either left-going or right-going off/on trials to be identified as a time cell.

### Split-half reliability of sequences and single units

For single units, a random set of trials was selected from laser-off and laser-on trials, and Pearson correlation coefficient was calculated using resulting mean firing vectors. For the same laser condition comparison (off vs. off, on vs. on), the second set of trials was randomly selected from left-out trials. This was repeated 5000 times, and the mean of each null distribution was obtained. For sequences, a random set of trials was selected following the same procedure as described above, but was done separately for left off/on and right off/on trials. Pearson correlation coefficient was calculated using concatenated mean firing vectors. For all analyses, a number of trials used to calculate mean firing rate vectors was matched.

### Visualization of neural trajectories

For visualization purposes only, principal component analysis was performed as previously described (Cunningham & Yu, 2014) to reduce the dimensionality of smoothed spike trains (bin size = 200ms, S.D. = 1s). First three components were extracted, and PCA scores were plotted in a 3D state space. Only those sessions with at least six trials in all four conditions (left-going off/on, right-going off/on) were included in this visualization.

### Maximum likelihood estimation of delay time

A naive Bayesian classifier (K. Zhang et al., 1998) was implemented to decode time of the treadmill delay period. This classifier was originally intended for place cells and for decoding spatial locations. Accordingly, the prior probability distribution could be constructed out of the spatial occupancy map. However, given that time is constant, this prior probability becomes uniform in our application, and the resulting approach is equivalent to *maximum likelihood estimation*. That is, the log-likelihood is maximized with respect to delay time:

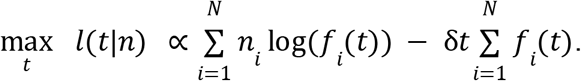

Here, for *N* neurons, the mean number of spikes fired by neuron *i* at delay time *t*is given by *f*_*i*_(*t*), and the delay time is decoded on the population spike vector *n*_*i*_. Spike trains were binned using a window size of *δt= 200*ms as with before and smoothed using a *σ* = 1s Gaussian kernel. Firing rates of zero (which normally leads to the undesirable condition of log(0) = − ∞) were replaced by a constant penalty term of 2^-52^ (eps in MATLAB). Due to the small number of simultaneously recorded time cells within a typical recording session, we pooled together neurons from all recordings that had at least 6 trials in each of four combinations of laser and behavioral choice conditions. An *m-out-of-n bootstrap* method (Bickel & Sakov, 2008) was subsequently used to estimate an empirical distribution of decoding accuracy, where *m = 50* neurons were sampled from the *n* total recorded neurons with replacement. This alternative *bootstrap* approach permits the number of neurons to be matched between the ArchT and the YFP groups to control for bias in sample sizes. The decoding accuracy was measured by *leave-one-out* cross-validation for each bootstrap sample by holding out single trials.

### Gini impurity index and decoding of behavioral choice

A custom classifier based on the *Gini impurity index* was used to decode animals’ choice (left or right trials) subsequent to the delay period, and hence to serve as a proxy measure of a neuron’s trajectory-dependency. This index is related to the measure of *entropy* in information theory and is frequently used for building *decision tree classifiers (Yang, 2010)*. Intuitively, if a neuron’s firing rate can be used to predict subsequent behavioral choice, then a certain threshold θ can be set so that trials in which the firing rate falls below the threshold are confined to one side of the maze, while the complement set of trials falls under the other side (**Figure S6**). In other words, by ordering the trials based on the neuron’s firing rate and splitting the trials by some appropriate threshold, a neuron that differentiates between left and right trials will have mostly left-going trials over one side of the threshold, while the other side will be predominantly right-going trials (**Figure S6**). The Gini index helps in establishing this threshold by quantify the “impurity” of a dataset:

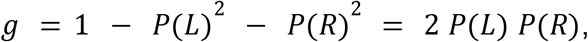

where *P*(*L*) and *P*(*R*) are the fractions of left-going and right-going trials respectively. The resulting measure ranges between 0 and 0.5, with 0 signifying that the data contains only one trial label (e.g., only left-going trials) and 0.5 defining an equally proportioned mixture.

First, a baseline Gini index *g*_0_ is computed from all trials in a given session. Then, the trials are sorted based on the firing rate of a neuron *f*_*k*_ (*t*) on each trial *k* at each time bin *t*, such that *f*_*k*_(*t*) ≤ *f*_*k* + 1_(*t*) for all *k* ∈ {1, 2, …, *K* − 1} over a total of *K* trials. The time series were smooth with a σ = 200ms Gaussian kernel to compensate for potential jitters in time fields. The trials can then be split into two subsets of trials at any of the *K* – 1 locations, in which case we can define the firing rate threshold 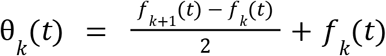 as the midway point between the two firing rates on those trials. After splitting the trials into two subsets (i.e., the upper and lower subsets), we compute the Gini impurity for each of them:

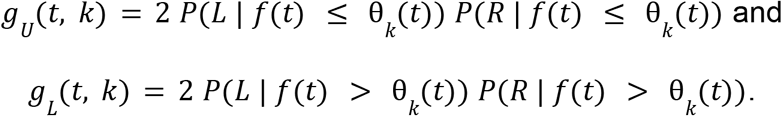

Finally, the Gini gain (i.e., the gain in purity) is calculated as the difference in impurity before and after splitting the trials, weighed by the size of the trial subsets:

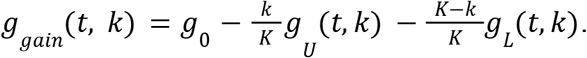

Effectively, a high Gini gain signifies that by placing the trials into two categories based on whether the neuron’s firing rate is lesser or greater than a threshold, left- and right-going trials can be accurately dissociated. Therefore, by maximizing Gini gain with respect to *t* and *k*, we estimate the time bin during the treadmill delay period and the firing rate threshold that gives the maximal accuracy in classifying behavioral choice. The decoding accuracy was assessed by a *leave-one-out* cross-validation approach.

### Identification of trajectory-dependent time cells

Trajectory-dependent time cells were identified following the procedure described previously (Kinsky et al., 2020). In short, the actual difference between mean firing rates of left-going and right-going trials was compared to the difference obtained by shuffling trial identity. The actual difference had to exceed the shuffled difference for 2850 out of 3000 times (95%). In order to avoid spurious detection of trajectory-dependent cells, cells had to have at least 3 consecutive bins that are significantly different, and at least 3 spikes in 40% of either left-going or right-going trials. Raw spike trains were used to ensure that these criteria are not simply met by smoothing. The identification was done separately for laser-off and on conditions.

### Histology

After mice were anesthetized deeply with isoflurane (5%), they were intracardially perfused with 1 x PBS and 4% paraformaldehyde. Brains were retrieved after animal heads were left in 4% paraformaldehyde for at least 48 hours. Subsequently, brains were kept in a 30% sucrose solution until they sank to the bottom. Following these procedures, brains were kept in a -80ºC freezer. Coronal sections were sliced using a cryostat at 20µm or 40µm. Both hippocampal and medial septum sections were mounted using a fluorescent DAPI labeling medium (Southern Biotechnology), and were imaged using a slide scanner (Olympus, VS120) or Zeiss Axio Observer to confirm tetrode tracks, optic fiber location, and viral vector expression. For YFP controls only, GFP amplification immunohistochemistry was performed. Sections were initially washed 3 × 5 minutes in 1 x PBS, and blocked 3 × 15 minutes in PGT solution (0.45% gelatin and 0.25% triton in 1 x PBS) under gentle agitation. Then, they were incubated in anti-GFP primary antibody (1:1000 dilution in PGT, anti-GFP rabbit (IgG), ThermoFisher, A-11122) overnight at 4°C (up to 24 hours). On the following day, they were washed 3 × 15 minutes in PGT solution and further incubated in Alexa Fluor-488 secondary antibody (1:1000 dilution in PGT, anti-rabbit goat Alexa Fluor-488 (IgG), ThermoFisher, A-21206) for 3 hours. They were washed 3 × 5 minutes in 1 x PBS, and mounted as described above. Histological data from mice #5653 (ArchT) and #7415 (YFP) could not be recovered due to misplacing their brains during storage, however in both cases the neurophysiology and response to laser were qualitatively similar to their respective groups and were included in the analysis.

### Statistical analysis

All statistical evaluations were performed under MATLAB and GNU R. Non-parametric Wilcoxon signed rank tests, Wilcoxon rank sum tests, Kolmogorov-Smirnov tests, and Kruskal-Wallis one-way ANOVAs were used throughout the paper. P-values reported from all *post hoc* tests were Bonferroni-adjusted for multiple comparisons. Robust statistics were performed using the ‘WRS’ package (Wilcox, 2021). Robust two-way mixed bootstrapped ANOVA was used to compare choice decoding accuracy across training-testing conditions between trajectory-dependent and trajectory-independent cells. Multiple paired bootstrap tests for equal mean was utilized to compare accuracy of time decoding between training-testing conditions. Statistical tests used for each figure were described in the corresponding figure panel. Box and whiskers represent IQR and 1.5 x IQR, respectively. SEM and 95% confidence interval were used throughout, and stated in each figure panel. Correlations were calculated using Pearson’s correlation coefficient.

## Acknowledgments

We graciously thank S. Kim, R. Lavoie, and V. Bazinet for technical assistance. We are also thankful to J. Ying, J.Q. Lee, M. Yaghoubi, J. Robinson, J.R. Hinman M.E. Hasselmo, and A.T. Keinath who provided comments and advice on prior versions of this manuscript and to all members of the Brandon laboratory for helpful discussions. H.C. is supported by the Canada Graduate Scholarships for Doctoral Program (NSERC CGS-D). This work was supported with funding from the Canadian Institutes for Health Research grants (Project grants #367017 and #377074), the Natural Sciences and Engineering Council (Discovery grant #74105), and the Canada Research Chairs Program to M.P.B.

## Author contributions

H.Y. and M.P.B. contributed to experimental design. H.Y. contributed to recordings. H.Y., H.C. and M.P.B. all contributed to analysis of data, and wrote the manuscript.

## Materials & Correspondence

Correspondence to Mark P. Brandon.

## Data availability

Source data for all experiments are publicly available at [insert Dryad link] or via request to the corresponding author.

## Code availability

All custom codes written for reported analyses are publicly available at [insert Github link] or via request to the corresponding author.

## Supplemental Information

**Supplemental figures S1-6**.

**Tables S1-4**

**Figure S1.**
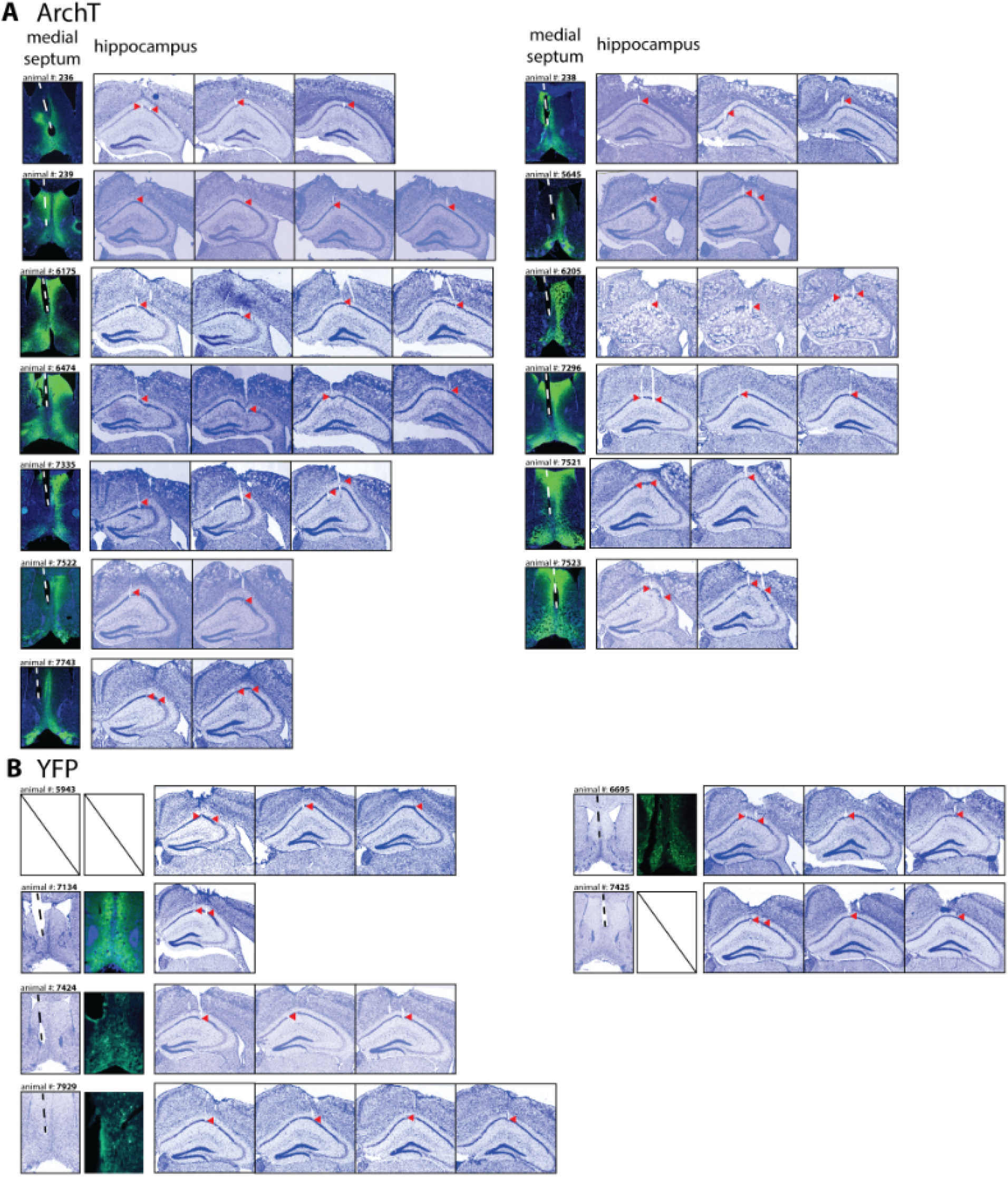
(A) Optic fiber track (white dashed line) with GFP expression in the MS for ArchT animals. Tetrode tracks (red arrows) in dorsal CA1 for ArchT animals. (B) Same as (A), but for YFP controls. Missing panels for YFP animals were due to damage during immunohistochemistry (#5943 / #7425).

**Figure S2.**
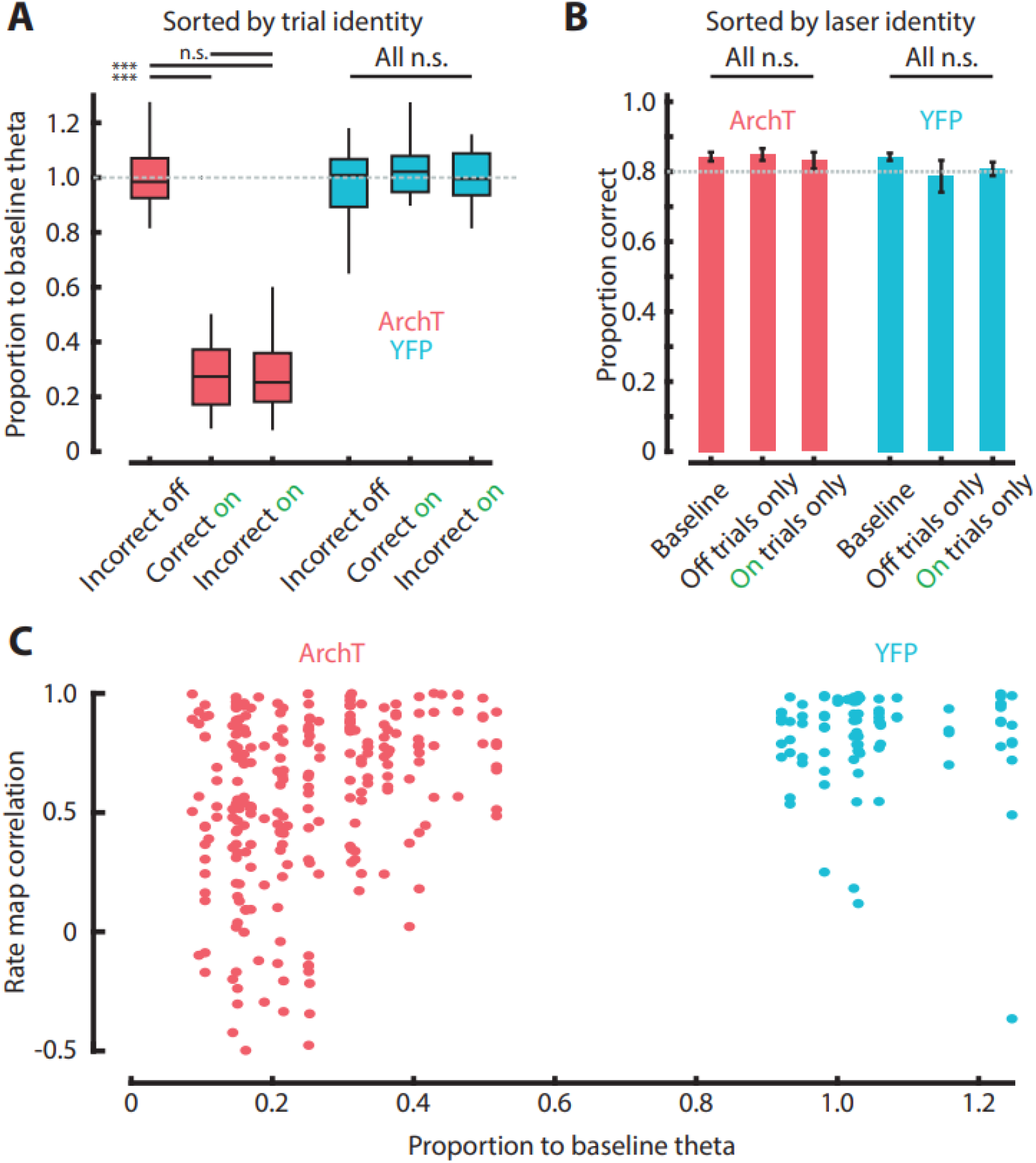
(A) Proportion to baseline theta as a function of correct and incorrect choice. There was no significant change in theta power between correct on and incorrect on trials (Kruskal-Wallis test followed by post hoc test, corrected for multiple comparisons with Bonferroni, ArchT: F(3,152) = 116.59, p = 4.18 × 10^−25^, YFP: F(3, 52) = 0.51, p = 0.92). Refer to Table S1 for post hoc test results. (B) Proportion correct based on laser identity. No behavior disruption was observed (Kruskal-Wallis test, ArchT: F(2, 39) = 0.61, p = 0.74, YFP: F(2,18) = 1.87, p = 0.39). (C) Pearson correlation between rate map correlation and proportion to baseline theta. Significant correlation observed in ArchT (r = 0.27, p = 1.65 × 10^−5^), but not in YFP (r = -0.05, p = 0.68).

**Figure S3.**
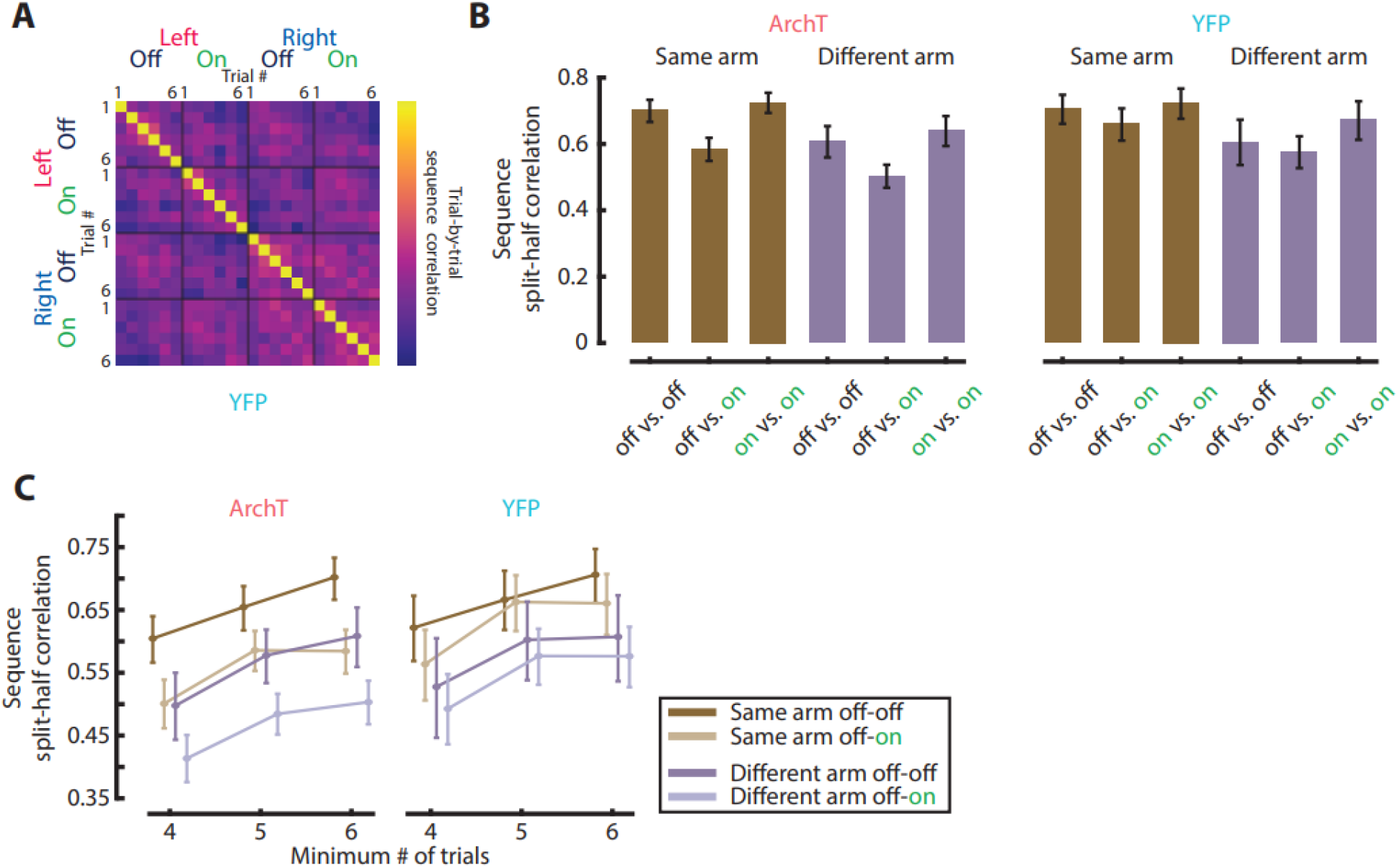
(A) Sequence similarity correlation matrix for YFP controls. (B) Split-half correlation for sequences. Trials were randomly divided into two parts, and the mean firing rate of these two parts was concatenated to build two sequences. Pearson correlation of these two sequences was obtained. This process was repeated 10,000 times. Bar plots indicate the mean and error bars indicate 95% confidence interval. Sessions with at least 6 trials of leff-going off/on and right-going off/on were used for this analysis. (C) Change in sequence split-half correlation as a function of minimum number of trials needed to be included in the analysis. Dots indicate the mean and error bars indicate 95% confidence interval. In all cases, cross-training condition (off-on, shaded color) was lower than its counterpart condition (off-off, solid color). Moreover, sequence split-half correlation for the same arm was higher than the different arm condition, suggesting that a checkered pattern in ArchT animals is observed regardless of threshold used for analysis.

**Figure S4.**
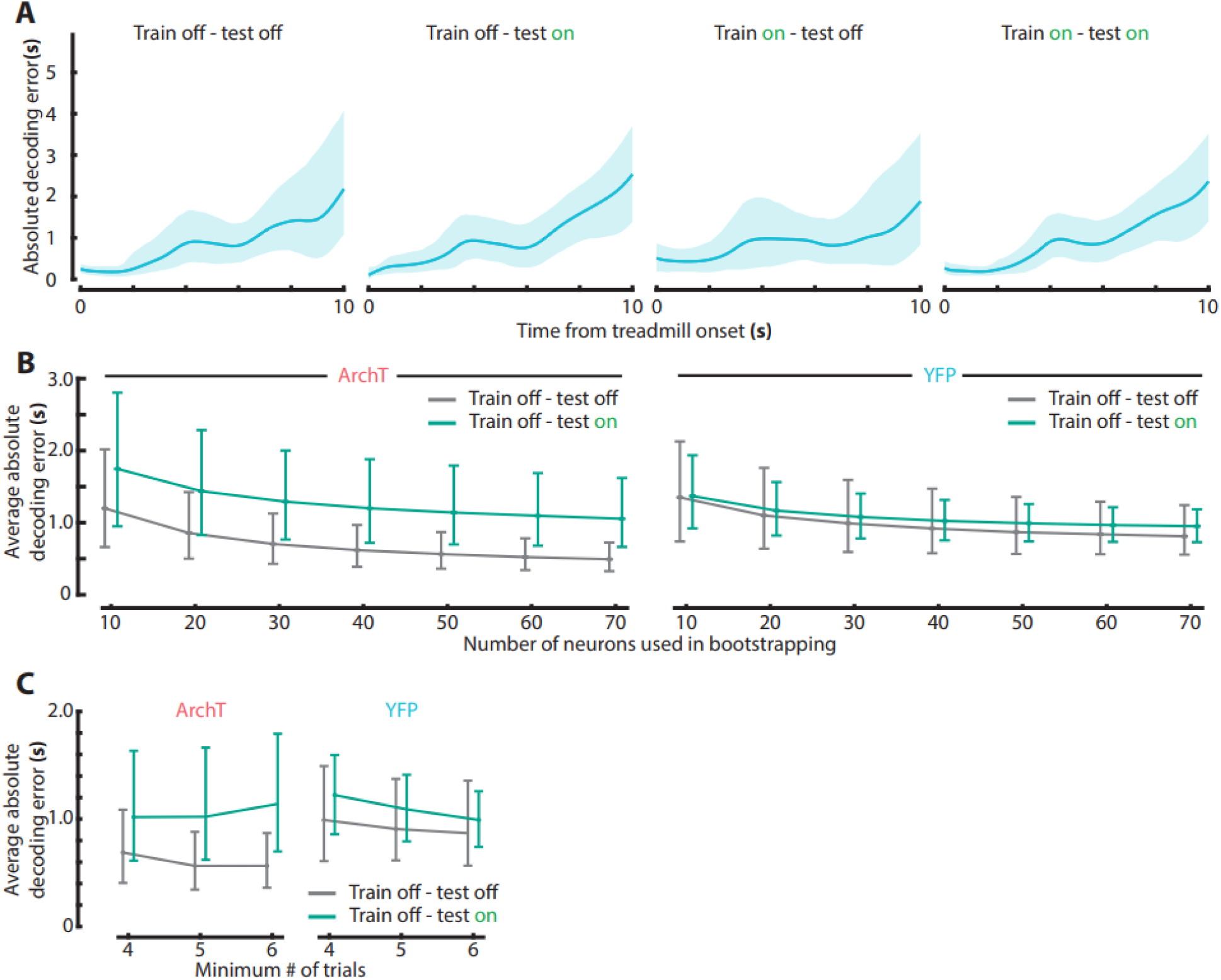
(A) Absolute decoding error as a function of time for YFP controls. (B) Average absolute decoding error as a function of the number of neurons used for bootstrapping. Note that off-off error (grey) is consistently lower than cross-training condition (off-on, green) for the ArchT group, but not for YFP controls. Dots indicate the mean and error bars indicate 95% confidence interval. (C) Average absolute decoding error as a function of minimum number of trials used. Similar to (B), ArchT animals have lower error in cross-training condition regardless of the minimum number of trials used to select sessions.

**Figure S5.**
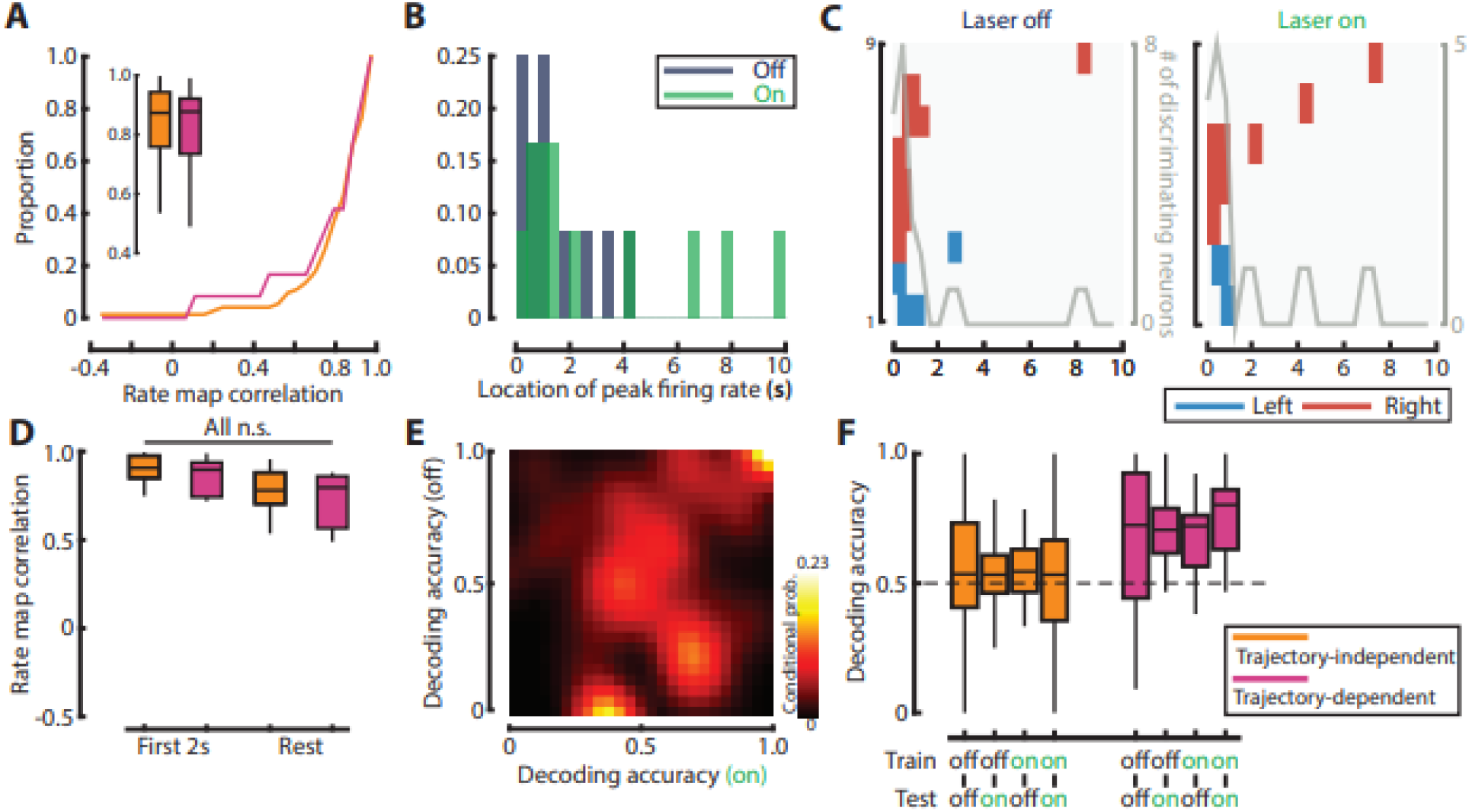
(A) Rate map correlation of YFP trajectory-independent and trajectory-dependent cells (Wilcoxon rank-sum test, YFP: p = 0.79). (B) Peak firing field location during laser off and on for YFP cells. (C) Significantly discriminating field locations for YFP. Note that they are concentrated at the beginning similar to ArchT cells. (D) Rate map correlation separated by the first 2 seconds vs. the rest. No significant difference observed. (E) Conditional probability matrix for decoding accuracy for YFP controls. (F) Decoding accuracy for trajectory-independent and trajectory dependent neurons. Refer to Table S3 for p values.

**Figure S6.**
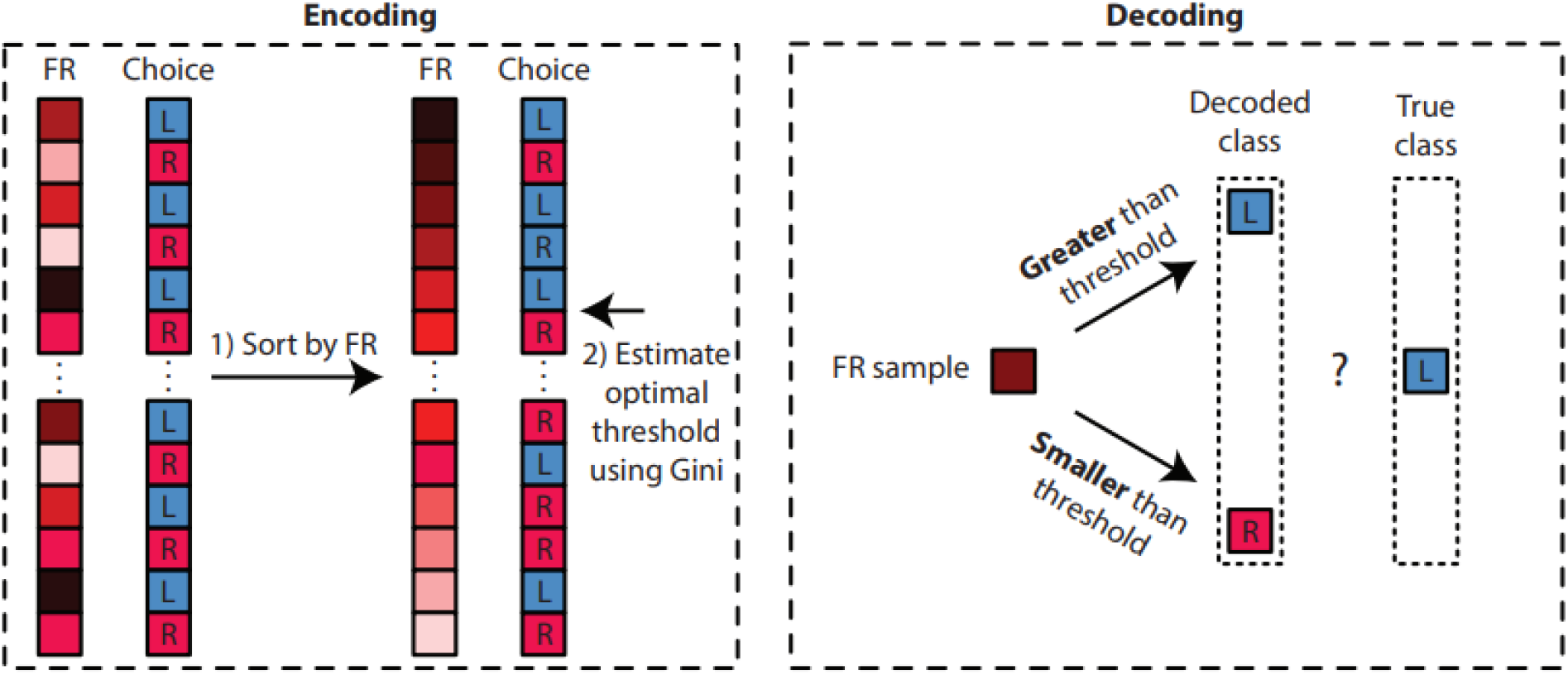
Overview of decoding method for animals’ choice following the treadmill run. Refer to methods for detailed description.

**Table S1.** post hoc p-values corrected for multiple comparisons with Bonferroni (Figure S2B).

| **ArchT** | Correct off | Correct on | Incorrect off | Incorrect on |
| --- | --- | --- | --- | --- |
| Correct off |  | **2.34 x 10<sup>-14</sup>** | 1.00 | **1.84 x 10<sup>-14</sup>** |
| Correct on |  |  | **1.05 x 10<sup>-12</sup>** | 1.00 |
| Incorrect off |  |  |  | **8.39 x 10<sup>-13</sup>** |

**Table S2.**
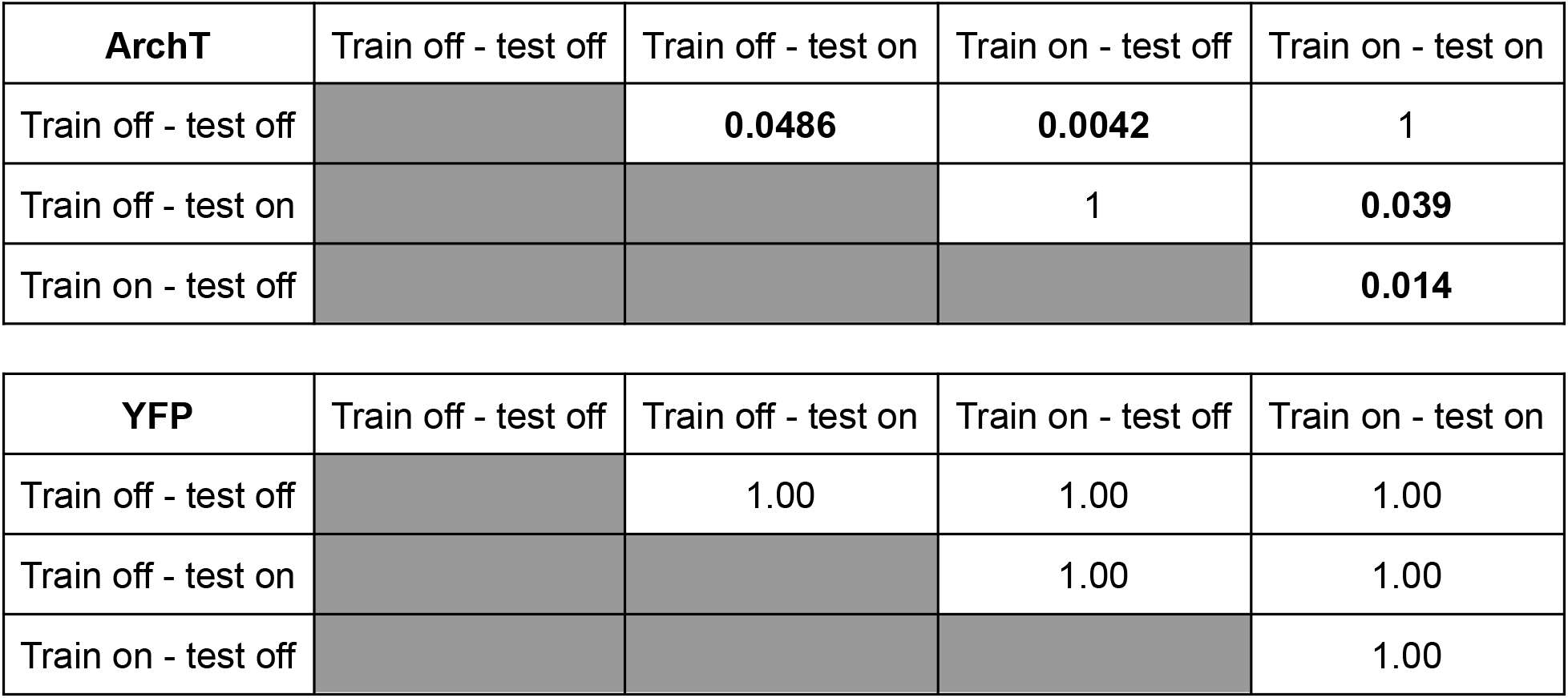
Paired-sample bootstrap for equal mean absolute error, Bonferroni corrected for multiple comparisons

| **ArchT** | Train off - test off | Train off - test on | Train on - test off | Train on - test on |
| --- | --- | --- | --- | --- |
| Train off - test off |  | **0.0486** | **0.0042** | 1 |
| Train off - test on |  |  | 1 | **0.039** |
| Train on - test off |  |  |  | **0.014** |

| **YFP** | Train off - test off | Train off - test on | Train on - test off | Train on - test on |
| --- | --- | --- | --- | --- |
| Train off - test off |  | 1.00 | 1.00 | 1.00 |
| Train off - test on |  |  | 1.00 | 1.00 |
| Train on - test off |  |  |  | 1.00 |

**Table S3.** Robust 2-way mixed bootstrapped ANOVA with 20% trimmed means

| **ArchT** |  |  |
| --- | --- | --- |
| **Type** | **Factor** | **P-value** |
| Between | Cell identity (Trajectory-dependent vs. independent) | **<1.00 x 10<sup>-4</sup>** |
| Within | Decoding condition (off-off, off-on, on-off, on-on) | 0.0644 |
| Interaction | Cell identity x decoding condition | 0.1560 |

**Table S3.** Robust 2-way mixed bootstrapped ANOVA with 20% trimmed means
| **YFP** |  |  |
| --- | --- | --- |
| **Type** | **Factor** | **P-value** |
| Between | Cell identity (Trajectory-dependent vs. independent) | **0.0062** |
| Within | Decoding condition (off-off, off-on, on-off, on-on) | 0.2008 |
| Interaction | Cell identity x decoding condition | 0.1764 |

**Table S4.** Average weight of female and male VGAT::Cre mice.

|  | Male | Female |
| --- | --- | --- |
| 8 weeks old<br>(n = 5 per group) | 23.8 ± 1.10 (SD) | 17.6 ± 0.89 (SD) |
| 16-17 weeks old<br>(n = 6 per group) | 28.8 ± 0.98 (SD) | 21.8 ± 1.17 (SD) |

## References

Ahmed, M. S., Priestley, J. B., Castro, A., Stefanini, F., Solis Canales, A. S., Balough, E. M., Lavoie, E., Mazzucato, L., Fusi, S., & Losonczy, A. (2020). Hippocampal Network Reorganization Underlies the Formation of a Temporal Association Memory. Neuron, 107(2), 283–291.e6.

Bickel, P. J., & Sakov, A. (2008). ON THE CHOICE OF m IN THE m OUT OF n BOOTSTRAP AND CONFIDENCE BOUNDS FOR EXTREMA. Statistica Sinica, 18(3), 967–985.

Bonett, D. G., & Wright, T. A. (2000). Sample size requirements for estimating pearson, kendall and spearman correlations. In Psychometrika (Vol. 65, Issue 1, pp. 23–28). https://doi.org/10.1007/bf02294183

Boyce, R., Glasgow, S. D., Williams, S., & Adamantidis, A. (2016). Causal evidence for the role of REM sleep theta rhythm in contextual memory consolidation. In Science (Vol. 352, Issue 6287, pp. 812–816). https://doi.org/10.1126/science.aad5252

Brandon, M. P., Bogaard, A. R., Libby, C. P., Connerney, M. A., Gupta, K., & Hasselmo, M. E. (2011). Reduction of theta rhythm dissociates grid cell spatial periodicity from directional tuning. Science, 332(6029), 595–599.

Brandon, M. P., Koenig, J., Leutgeb, J. K., & Leutgeb, S. (2014). New and Distinct Hippocampal Place Codes Are Generated in a New Environment during Septal Inactivation. In Neuron (Vol. 82, Issue 4, pp. 789–796). https://doi.org/10.1016/j.neuron.2014.04.013

Brun, V. H., Leutgeb, S., Wu, H.-Q., Schwarcz, R., Witter, M. P., Moser, E. I., & Moser, M.-B. (2008). Impaired spatial representation in CA1 after lesion of direct input from entorhinal cortex. Neuron, 57(2), 290–302.

Brun, V. H., Otnass, M. K., Molden, S., Steffenach, H.-A., Witter, M. P., Moser, M.-B., & Moser, E. I. (2002). Place cells and place recognition maintained by direct entorhinal-hippocampal circuitry. Science, 296(5576), 2243–2246.

Buzsáki, G., & Moser, E. I. (2013). Memory, navigation and theta rhythm in the hippocampal-entorhinal system. Nature Neuroscience, 16(2), 130–138.

Cunningham, J. P., & Yu, B. M. (2014). Dimensionality reduction for large-scale neural recordings. Nature Neuroscience, 17(11), 1500–1509.

Driscoll, L. N., Pettit, N. L., Minderer, M., Chettih, S. N., & Harvey, C. D. (2017). Dynamic Reorganization of Neuronal Activity Patterns in Parietal Cortex. Cell, 170(5), 986–999.e16.

Eichenbaum, H. (2017). On the Integration of Space, Time, and Memory. In Neuron (Vol. 95, Issue 5, pp. 1007–1018). https://doi.org/10.1016/j.neuron.2017.06.036

Etter, G., van der Veldt, S., Choi, J., & Williams, S. (2021). Optogenetic frequency scrambling of hippocampal theta oscillations dissociates working memory retrieval from hippocampal spatiotemporal codes. bioRxiv. https://doi.org/10.1101/2021.12.24.474139

Frank, L. M., Brown, E. N., & Wilson, M. (2000). Trajectory encoding in the hippocampus and entorhinal cortex. Neuron, 27(1), 169–178.

Gemzik, Z. M., Donahue, M. M., & Griffin, A. L. (2021). Optogenetic suppression of the medial septum impairs working memory maintenance. Learning & Memory, 28(10), 361–370.

Gill, P. R., Mizumori, S. J. Y., & Smith, D. M. (2011). Hippocampal episode fields develop with learning. Hippocampus, 21(11), 1240–1249.

Haimerl, C., Angulo-Garcia, D., Villette, V., Reichinnek, S., Torcini, A., Cossart, R., & Malvache, A. (2019). Internal representation of hippocampal neuronal population spans a time-distance continuum. Proceedings of the National Academy of Sciences of the United States of America, 116(15), 7477–7482.

Han, X., Chow, B. Y., Zhou, H., Klapoetke, N. C., Chuong, A., Rajimehr, R., Yang, A., Baratta, M. V., Winkle, J., Desimone, R., & Boyden, E. S. (2011). A High-Light Sensitivity Optical Neural Silencer: Development and Application to Optogenetic Control of Non-Human Primate Cortex. In Frontiers in Systems Neuroscience (Vol. 5). https://doi.org/10.3389/fnsys.2011.00018

Hasselmo, M. E. (1999). Neuromodulation: acetylcholine and memory consolidation. Trends in Cognitive Sciences, 3(9), 351–359.

Hasselmo, M. E. (2008). Grid cell mechanisms and function: contributions of entorhinal persistent spiking and phase resetting. Hippocampus, 18(12), 1213–1229.

Hasselmo, M. E., & Stern, C. E. (2014). Theta rhythm and the encoding and retrieval of space and time. NeuroImage, 85 Pt 2, 656–666.

Kinsky, N. R., Mau, W., Sullivan, D. W., Levy, S. J., Ruesch, E. A., & Hasselmo, M. E. (2020). Trajectory-modulated hippocampal neurons persist throughout memory-guided navigation. Nature Communications, 11(1), 2443.

Koenig, J., Linder, A. N., Leutgeb, J. K., & Leutgeb, S. (2011). The Spatial Periodicity of Grid Cells Is Not Sustained During Reduced Theta Oscillations. In Science (Vol. 332, Issue 6029, pp. 592–595). https://doi.org/10.1126/science.1201685

Kraus, B. J., Brandon, M. P., Robinson, R. J., 2nd, Connerney, M. A., Hasselmo, M. E., & Eichenbaum, H. (2015). During Running in Place, Grid Cells Integrate Elapsed Time and Distance Run. Neuron, 88(3), 578–589.

Kraus, B. J., Robinson, R. J., 2nd, White, J. A., Eichenbaum, H., & Hasselmo, M. E. (2013). Hippocampal “time cells”: time versus path integration. Neuron, 78(6), 1090–1101.

Levy, S. J., Kinsky, N. R., Mau, W., Sullivan, D. W., & Hasselmo, M. E. (2021). Hippocampal spatial memory representations in mice are heterogeneously stable. Hippocampus, 31(3), 244–260.

Li, N., Chen, S., Guo, Z. V., Chen, H., Huo, Y., Inagaki, H. K., Chen, G., Davis, C., Hansel, D., Guo, C., & Svoboda, K. (2019). Spatiotemporal constraints on optogenetic inactivation in cortical circuits. eLife, 8. https://doi.org/10.7554/eLife.48622

MacDonald, C. J., Carrow, S., Place, R., & Eichenbaum, H. (2013). Distinct hippocampal time cell sequences represent odor memories in immobilized rats. The Journal of Neuroscience: The Official Journal of the Society for Neuroscience, 33(36), 14607–14616.

MacDonald, C. J., Lepage, K. Q., Eden, U. T., & Eichenbaum, H. (2011). Hippocampal “time cells” bridge the gap in memory for discontiguous events. Neuron, 71(4), 737–749.

MacDonald, C. J., & Tonegawa, S. (2021). Crucial role for CA2 inputs in the sequential organization of CA1 time cells supporting memory. In Proceedings of the National Academy of Sciences (Vol. 118, Issue 3). https://doi.org/10.1073/pnas.2020698118

Maldonado, M. A., Allred, R. P., Felthauser, E. L., & Jones, T. A. (2008). Motor skill training, but not voluntary exercise, improves skilled reaching after unilateral ischemic lesions of the sensorimotor cortex in rats. Neurorehabilitation and Neural Repair, 22(3), 250–261.

Marr, D. (1971). Simple Memory: A Theory for Archicortex. Philos Trans R Soc Lond B Biol Sci, 262(841), 23–81.

Mau, W., Sullivan, D. W., Kinsky, N. R., Hasselmo, M. E., Howard, M. W., & Eichenbaum, H. (2018). The Same Hippocampal CA1 Population Simultaneously Codes Temporal Information over Multiple Timescales. Current Biology: CB, 28(10), 1499–1508.e4.

McNaughton, B. L., & Morris, R. G. M. (1987). Hippocampal synaptic enhancement and information storage within a distributed memory system. Trends in Neurosciences, 10(10), 408–415.

Mitra, P., & Bokil, H. (2007). Observed Brain Dynamics. https://doi.org/10.1093/acprof:oso/9780195178081.001.0001

Otchy, T. M., Wolff, S. B. E., Rhee, J. Y., Pehlevan, C., Kawai, R., Kempf, A., Gobes, S. M. H., & Ölveczky, B. P. (2015). Acute off-target effects of neural circuit manipulations. Nature, 528(7582), 358–363.

Pastalkova, E., Itskov, V., Amarasingham, A., & Buzsáki, G. (2008). Internally generated cell assembly sequences in the rat hippocampus. Science, 321(5894), 1322–1327.

Robinson, N. T. M., Priestley, J. B., Rueckemann, J. W., Garcia, A. D., Smeglin, V. A., Marino, F. A., & Eichenbaum, H. (2017). Medial Entorhinal Cortex Selectively Supports Temporal Coding by Hippocampal Neurons. Neuron, 94(3), 677–688.e6.

Roland, J. J., Janke, K. L., Servatius, R. J., & Pang, K. C. H. (2014). GABAergic neurons in the medial septum-diagonal band of Broca (MSDB) are important for acquisition of the classically conditioned eyeblink response. Brain Structure & Function, 219(4), 1231–1237.

Sabariego, M., Schönwald, A., Boublil, B. L., Zimmerman, D. T., Ahmadi, S., Gonzalez, N., Leibold, C., Clark, R. E., Leutgeb, J. K., & Leutgeb, S. (2019). Time Cells in the Hippocampus Are Neither Dependent on Medial Entorhinal Cortex Inputs nor Necessary for Spatial Working Memory. Neuron, 102(6), 1235–1248.e5.

Salz, D. M., Tiganj, Z., Khasnabish, S., Kohley, A., Sheehan, D., Howard, M. W., & Eichenbaum, H. (2016). Time Cells in Hippocampal Area CA3. In Journal of Neuroscience (Vol. 36, Issue 28, pp. 7476–7484). https://doi.org/10.1523/jneurosci.0087-16.2016

Shimbo, A., Izawa, E.-I., & Fujisawa, S. (2021). Scalable representation of time in the hippocampus. Science Advances, 7(6). https://doi.org/10.1126/sciadv.abd7013

Skaggs, W. E., McNaughton, B. L., Wilson, M. A., & Barnes, C. A. (1996). Theta phase precession in hippocampal neuronal populations and the compression of temporal sequences. Hippocampus, 6(2), 149–172.

Tulving, E. (1983). Elements of Episodic Memory. Oxford University Press, USA.

Wang, Y., Romani, S., Lustig, B., Leonardo, A., & Pastalkova, E. (2015). Theta sequences are essential for internally generated hippocampal firing fields. Nature Neuroscience, 18(2), 282–288.

Wilcox, R. R. (2021). Introduction to Robust Estimation and Hypothesis Testing. Academic Press.

Wood, E. R., Dudchenko, P. A., Robitsek, R. J., & Eichenbaum, H. (2000). Hippocampal neurons encode information about different types of memory episodes occurring in the same location. Neuron, 27(3), 623–633.

Yang, Z. R. (2010). Machine Learning Approaches to Bioinformatics. World Scientific.

Zhang, K., Ginzburg, I., McNaughton, B. L., & Sejnowski, T. J. (1998). Interpreting neuronal population activity by reconstruction: unified framework with application to hippocampal place cells. Journal of Neurophysiology, 79(2), 1017–1044.

Zhang, Y., Cao, L., Varga, V., Jing, M., Karadas, M., Li, Y., & Buzsáki, G. (2021). Cholinergic suppression of hippocampal sharp-wave ripples impairs working memory. Proceedings of the National Academy of Sciences of the United States of America, 118(15). https://doi.org/10.1073/pnas.2016432118

Ziv, Y., Burns, L. D., Cocker, E. D., Hamel, E. O., Ghosh, K. K., Kitch, L. J., El Gamal, A., & Schnitzer, M. J. (2013). Long-term dynamics of CA1 hippocampal place codes. Nature Neuroscience, 16(3), 264–266.

